# GDNF enemas improve epithelial and immune defects in both aganglionic and ganglionic colon of Hirschsprung mice

**DOI:** 10.64898/2026.08.31.748309

**Authors:** Nejia Lassoued, Jacob Trudel, Marie A Lefèvre, Alassane Gary, Zhiqiang Guo, Alexis Yero, Mohammad-Ali Jenabian, Rodolphe Soret, Nicolas Pilon

## Abstract

Hirschsprung disease (HSCR) is a severe birth defect where ganglia of the enteric nervous system (ENS) are missing from distal bowel. The aganglionic segment is also characterized by increased epithelial permeability and pro-inflammatory immune activation. These problems may sequentially lead to translocation of gut microbes into the colon wall and systemic circulation, resulting in enterocolitis and sepsis. Current HSCR treatment via surgical resection of the aganglionic segment is lifesaving but not curative, often leaving patients with persistent gastrointestinal complications including recurrent risk of enterocolitis. As alternative, we are developing a regenerative medicine strategy based on *in situ* stimulation of tissue-resident ENS progenitors via rectal administration of the neurotrophic factor GDNF. Here, we report that GDNF-based therapy has pleiotropic gastrointestinal effects in a mouse model of short-segment HSCR, beyond its role in ENS regeneration. Interestingly, we found that these protective effects are not restricted to the aganglionic distal colon, also positively impacting the ENS-containing proximal colon. GDNF treatment reduces bacterial translocation both locally and in peripheral organs, and this is associated with recovery of the key epithelial junction proteins CLDN3, ZO1 and DSG2. Furthermore, multiparameter flow cytometry-based analysis of 55 lymphoid and 17 myeloid cell subtypes revealed that GDNF treatment has global anti-inflammatory effects, preferentially affecting innate over adaptive immunity. Overall, these findings highlight a critical role for GDNF treatment in reestablishing proper epithelial and immune cell homeostasis, offering promising therapeutic avenues not only for HSCR but also potentially for other intestinal disorders with overlapping pathophysiology.

## INTRODUCTION

Hirschsprung disease (HSCR) is a congenital disorder of the enteric nervous system (ENS) characterized by the absence of enteric ganglia in the rectum and over variable lengths of the upstream bowel – being restricted to the distal colon in the most common short-segment form. Clinical presentation includes intestinal obstruction, abdominal distension, bilious vomiting and life-threatening complications such as Hirschsprung-associated enterocolitis (HAEC) [1, 2]. HSCR results from incomplete colonization of the bowel by neural crest-derived ENS progenitors, leaving the affected segment with a permanent lack of ENS ganglia [3–5]. While surgical resection of this so-called aganglionic segment remains the standard of care for HSCR, many patients continue to experience persistent gastrointestinal dysfunction and recurrent HAEC episodes after surgery [6–9].

Recent studies by our team have suggested that glial cell line-derived neurotrophic factor (GDNF), a critical regulator of ENS development, could be used for non-surgical treatment of HSCR [10, 11]. During prenatal development, canonical GDNF signaling through RET and GFRα1 receptors promotes ENS progenitor migration, proliferation, survival and differentiation in enteric neurons [12–14]. Yet, in the postnatal colon of short-segment HSCR mouse models, rectally administered GDNF has been shown to instead use NCAM1, not RET, as signaling receptor for inducing *de novo* enteric neurogenesis from various subtypes of tissue-resident ENS progenitors [10, 15]. Importantly, this intrarectal GDNF treatment was not only shown to restore functional ENS-like networks, but also to improve overall colon structure, microbiota composition and life expectancy of HSCR mice [5, 10, 11]. These findings suggest that GDNF treatment has broader effects in the colon than initially thought, including improvement/normalization of the intestinal epithelial barrier and immune system, but this hypothesis has not been fully explored in previous work.

The intestinal epithelial barrier represents a critical line of defense against potentially harmful luminal microbes, while allowing nutrients to pass through. Such selective permeability of the epithelial cell layer is mainly ensured by an array of intercellular junctional complexes, including tight junctions, adherens junctions, and desmosomes. Key components such as claudins, occludins, and zonula occludens proteins regulate paracellular permeability [16–20], while E-cadherin/catenins and desmogleins/desmocollins ensure adhesion and mechanical stability [21–24]. Disruption of these structures can lead to increased epithelial permeability, microbial translocation, and dysregulated pro-inflammatory immune activation [21, 25, 26]. All these problems are hallmarks of HSCR and are thought to underlie susceptibility to HAEC. Interestingly, GDNF has been previously reported to enhance the expression and organization of junctional proteins in mouse models of inflammatory bowel disease (IBD), thereby reinforcing epithelial barrier integrity and attenuating inflammatory responses [27–30]. This effect could be direct, at least in part, as some epithelial cell subsets including LGR5^+^ stem cells are known to express RET after birth [30–35].

Neuroimmune interactions also play a pivotal role in maintaining intestinal homeostasis, with ENS-derived signals modulating both innate and adaptive immune responses [36–40]. In the context of HSCR, the absence of enteric neurons disrupts this communication, contributing to exaggerated inflammation and increased risk of HAEC [10, 41]. Immune cell infiltration has been reported in HSCR mouse models, but previous studies consisted mostly in histological analyses without comprehensive immunophenotypic characterization [10, 42, 43]. Interestingly, GDNF has also been proposed to act directly on immune cells, as RET and/or NCAM1 are expressed in a large array of immune cell subsets [44–47]. Thus, a better understanding of how GDNF influences both epithelial barrier integrity and immune homeostasis in the treatment of HSCR is warranted.

This study provides new mechanistic insights into the pleiotropic effects of GDNF treatment on epithelial barrier function and immune regulation in homozygous *Holstein* (*Hol^Tg/Tg^*) mice, a well-established model of short-segment HSCR [48, 49]. Immunofluorescence and fluorescence *in situ* hybridization (FISH) were used to assess junctional proteins expression and bacterial translocation, while multicolor flow cytometry enabled detailed profiling of lymphoid and myeloid populations in colonic tissues. Our findings demonstrate that GDNF-based therapy partially restores epithelial barrier integrity and rebalance the immune populations through its anti-inflammatory effect, establishing a revised framework for understanding how GDNF treatment may globally heal the colon in the context of HSCR.

## MATERIALS AND METHODS

### Mice

Wild-type FVB mice (WT FVB/NCrl; Strain code 207) were purchased from Charles River Laboratories. *Hol^Tg/Tg^* mice were maintained on the FVB background as previously described [48]. Genotyping was performed by visual inspection of coat color [48]. All mice were housed in individually ventilated cages under 12 h light–12 h dark cycle (7 AM to 7 PM) with *ad libitum* access to standard chow (Charles River Rodent Diet #5075, Cargill Animal Nutrition). Where indicated, *Hol^Tg/Tg^*pups received daily enemas of 10 μl containing 1 μg/μl recombinant human GDNF (PeproTech #450-10, Cranbury, NJ) in phosphate-buffered saline (PBS) from postnatal day (P) 4 to P8, following our previous protocol [10] (Figure 1). Mice were euthanized at postnatal day (P) 20 by CO_2_ inhalation under isoflurane anesthesia. Both sexes were included in all experiments. Procedures complied with the Canadian Council on Animal Care (CCAC) guidelines and were approved by the Comité institutionnel de protection des animaux at UQAM (CIPA; reference #959).

**Figure 1.**
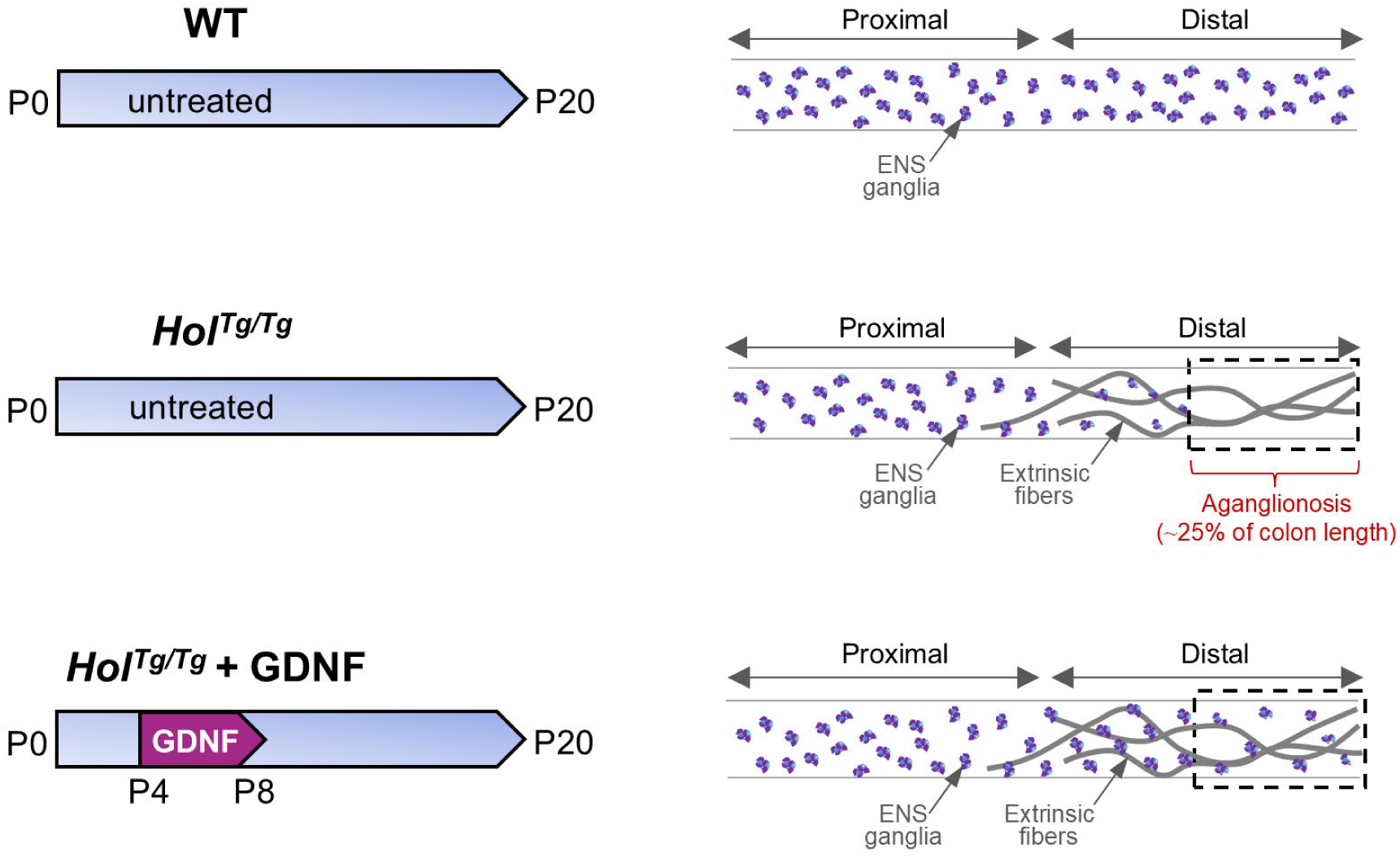
Schematic overview of experimental groups. For each group, the colon was dissected at P20 and split into two halves to analyze the proximal and distal segments separately. The colon of untreated WT mice is fully covered by ENS ganglia (upper panels), which are absent from the most distal 25% of the colon in untreated *Hol^Tg/Tg^* mice (middle panels). Hence, both this aganglionic region (highlighted by the dashed box) and upstream hypoganglionic transition zone are included in distal colon samples from *Hol^Tg/Tg^*mice. The distal colon of *Hol^Tg/Tg^* mice is also characterized by the presence of thick extrinsic nerve fibers. In *Hol^Tg/Tg^*mice that were previously treated with GDNF enemas between P4 to P8 (lower panels), ENS ganglia are regenerated in the otherwise aganglionic and hypoganglionic regions.

### Assessment of bacterial translocation

To evaluate systemic bacterial dissemination, peritoneal lavage fluid, liver, spleen and kidneys were collected from P20 mice, at which point most untreated *Hol^Tg/Tg^* mice are moribund. The peritoneal cavity was flushed with 1 ml sterile PBS, gently massaged, and aspirated to recover 400 μl of lavage fluid. Organs were weighed and homogenized in 400 μl of PBS using a tissues grinder (VWR). Aliquots of 100 μl of each sample (tissue homogenate or lavage fluid) were plated on gut microbiota medium (Anaerobe Systems, Inc, cat# AS-555) and chocolate agar (Thermo Fisher Scientific, cat# R01293) plates. For each sample, two replicate plates were prepared per medium and incubated under aerobic and anaerobic conditions, both at 37°C for 1-5 days. The anaerobic conditions were obtained by using an anaerobic jar (cat# 323-246-385) and anaerobic gas generator (cat# 323-246-376) from Thermo Fisher Scientific. Bacterial colonies were manually counted, and bacterial load was expressed in Colony-Forming Unit (CFU) per mg of tissue or ml of liquid. To assess potential contamination, negative controls were systematically included in each experiment by incubating culture plates containing sterile medium only. These control plates were processed exactly as the experimental plates, but yielded no colonies.

### Tissue collection and processing for immunofluorescence and FISH

The colon was dissected out from freshly euthanized mice in 1X PBS, flushed to remove fecal pellets, and divided into proximal and distal segments (Figure 1). These colon segments were then fixed in 4% paraformaldehyde (PFA) for 1 h at room temperature (RT) or overnight (ON) at 4 °C, rinsed three times with 1X PBS, and incubated ON in 30% sucrose. Tissues were finally embedded in OCT (Optimal Cutting Temperature) compound and cryosectioned at 15 μm (Leica CM1950). Sections were stored at -80 °C until processing.

For immunofluorescence or FISH, cryosections were thawed and washed three times in 1X PBS to remove residual OCT. Antigen retrieval was performed by incubating slides in retrieval buffer (Dako, Agilent, code# S2369) at 80°C for 35 min in a rice cooker, followed by cooling at RT for 20 min and three PBS washes. For immunofluorescence, sections were permeabilized and blocked for 1 h at RT in PBS containing 10% FBS and 1% Triton X100, then incubated with primary antibodies ON at 4°C and corresponding secondary antibodies for 2 h at RT. All antibodies and dilution factors are listed in Table S1. Nuclei were counterstained with DAPI (4’,6-diamidino-2-phenylindole) for 15 min at 1ug/ml. Slides were mounted in 100% glycerol and sealed with nail polish. FISH was performed to detect bacterial colonization using the Cy3-labeled Eub338 probe targeting bacterial 16S rRNA, and a Cy3-labeled negative control probe (Table S1), following established protocol [50].

### Imaging and data analysis

Images were acquired using a Nikon A1 confocal microscope (Melville, NY) using 20x or 60x objectives. Between 3 and 5 representative fields were captured per biological replicate. Image analysis was performed with ImageJ software. Background fluorescence was reduced using the “subtract background” function, while cell and bacterial counts were conducted with the “multi-point” tool. Fluorescence intensity measurements were calculated using the “Analyze” function.

### Tissue collection and processing for flow cytometry

Full-thickness proximal and distal colon tissues were dissected, cleaned, and mechanically dissociated in RPMI 1640 (Wisent inc, #350-000-CL) as previously described [51]. The resulting cell suspensions (600,000–1,000,000 viable cells each, in 200 μl), were stained with two antibody panels targeting lymphoid and myeloid markers following our established protocol [51]. Flow cytometry analysis was performed on an LSR Fortessa™ X-20 (BD Biosciences) with a 3-laser configuration (blue, violet, yellow-green). Cell populations were gated and analyzed using the FlowJo software (V10.10.0, BD Biosciences). All gating details can be found in our previous study [51].

### Statistical analysis

All experiments were conducted with at least 3 independent biological replicates. The exact number of replicates (n) and statistical tests used are specified in figures and/or accompanying legends. Data are presented as mean ± standard error of the mean (SEM). Statistical analyses were performed using GraphPad Prism (V10.6.1). Differences were considered significant at *p* < 0.05.

## RESULTS

### GDNF treatment limits bacterial translocation and septicemia in *Hol^Tg/Tg^* mice

Like most HSCR mouse models, *Hol^Tg/Tg^* mice maintained on the FVB background die around weaning age from complications of aganglionic megacolon [48]. The aganglionic segment of these mice is restricted to the distal colon, spanning ∼25% of total colon length (Figure 1), without sex bias [48]. To confirm that bacterial translocation and subsequent systemic infection contribute to their death, we first cultured liver, spleen, kidney, and peritoneal lavage samples from P20 mice (*i.e*., just before weaning) under both aerobic and anaerobic conditions, comparing WT, untreated *Hol^Tg/Tg^*and GDNF-treated *Hol^Tg/Tg^* animals (Figure 1). Under aerobic conditions on gut microbiota medium plates, samples from untreated *Hol^Tg/Tg^* mice exhibit a markedly elevated bacterial load in peripheral organs and peritoneal lavage, with levels ranging from 500 to 2500 CFU/g or ml (Figure 2a). A similar trend is observed under anaerobic conditions, where the bacterial load appears even more pronounced, reaching >3000 CFU/g in liver homogenates (Figure 2b). In stark contrast, *Hol^Tg/Tg^*mice that were previously treated with GDNF between P4-P8 display bacterial loads closer to WT levels (Figure 2a,b). Similar results were obtained when evaluating bacterial growth on chocolate agar plates (Figure S1a,b).

**Figure 2.**
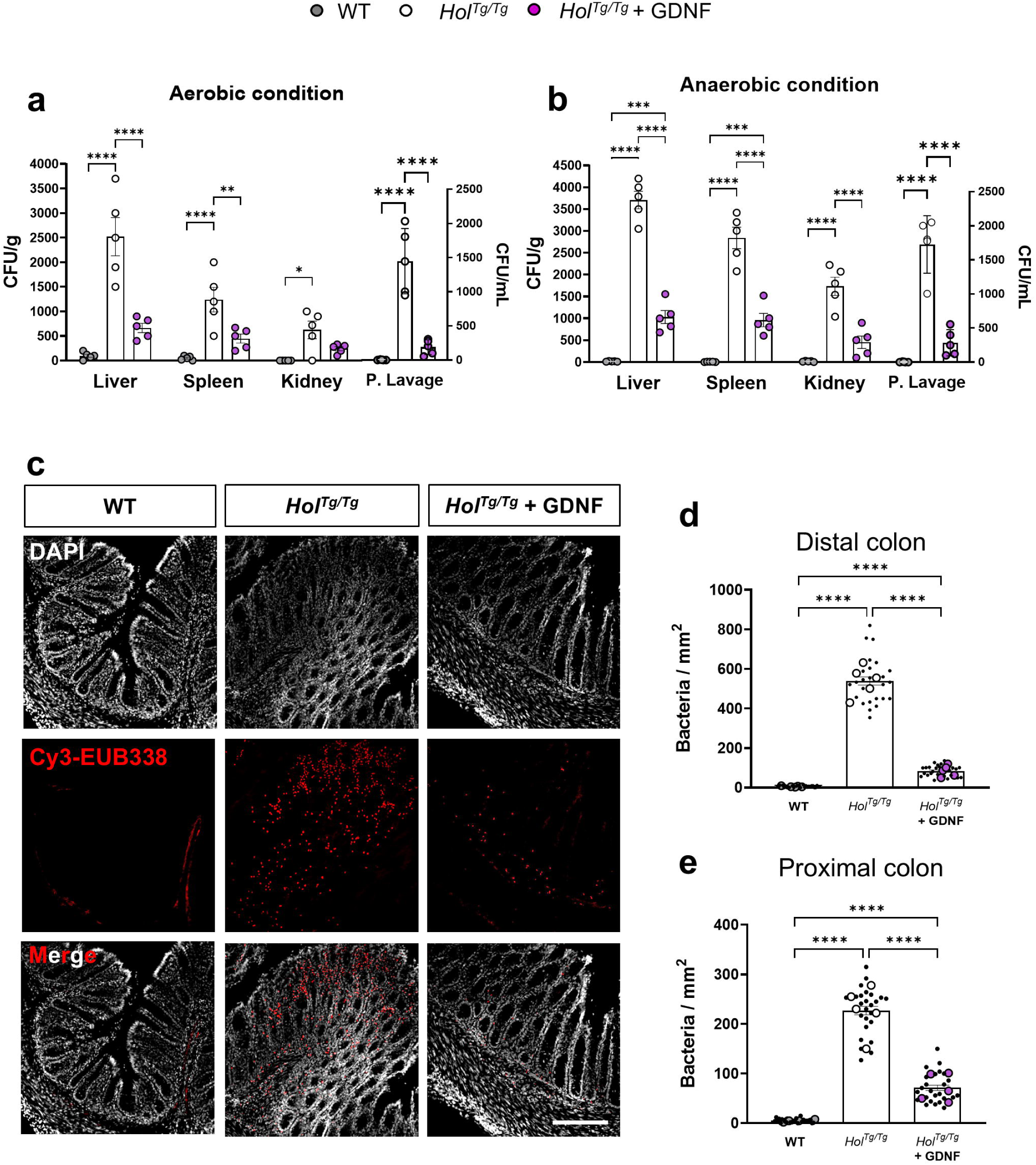
GDNF treatment reduces both local and peripheral bacterial translocation in P20 *Hol^Tg/Tg^*mice. (**a, b**) Quantitative analysis of bacterial load in samples of peritoneal lavage and peripheral organs (liver, spleen and kidneys) that were cultured on gut microbiota medium under aerobic (a) and anaerobic (b) conditions. GDNF treatment decreases back to WT-like levels the otherwise elevated bacterial load in samples from *Hol^Tg/Tg^* mice (n=5 mice per group). (**c**-**e**) FISH staining of bacterial 16S rRNA showing that GDNF treatment limits bacterial infiltration into the colonic mucosa of both distal (c,d) and proximal (e) colon from *Hol^Tg/Tg^* mice. Representative images in panel c correspond to 15µm-thick z-stack projections of distal colon samples (scale bar, 70 µm). Representative images of proximal colon samples can be found in Fig.S1. In the accompanying quantitative analyses shown in panels d-e, bacterial density is expressed as the number of bacteria per mm². Large dots represent the mean value for each mouse (n=5 mice per group), while small black dots correspond to each microscopic field analyzed (5 fields of view per animal). *p < .05, **p < .01, ***p < .001, ****p < .0001; Two-way (a,b) or one-way (d,e) ANOVA with Tukey’s post-hoc test.

To further substantiate these septicemia data, we also quantified local bacteria translocation in the mucosa layer of colon cross-sections from the same P20 mice using FISH. Consistent with the septicemia data, samples from *Hol^Tg/Tg^* mice display substantial bacterial colonization of the mucosa compared to WT controls (Figures 2c-e and S1c), in both proximal and distal halves of the colon (Figure 1). However, this alteration is more pronounced in the distal segment containing the aganglionic zone (Figure 1), where bacterial infiltration is >2-fold higher than in the ENS-containing proximal segment (Figure 2d,e) (227 ± 48 bacteria/mm^2^ in proximal colon vs 539 ± 77 bacteria/mm^2^ in distal colon). Remarkably, prior enema treatment with GDNF reduces mucosal bacterial density by a factor of 3-6X, reaching similar levels in both proximal (71 ± 27 bacteria/mm^2^) and distal colon (84 ± 28 bacteria/mm^2^) (Figure 2d,e). Collectively these data highlight the capacity of GDNF treatment to efficiently limit bacterial dissemination from the bowel to other sites.

### GDNF treatment improves epithelial barrier integrity and neutrophil counts in the colon of *Hol^Tg/Tg^* mice

To examine epithelial barrier integrity, colon cross-sections of P20 mice were also immunostained for key junctional proteins including the tight junction components Claudin-3 (CLDN3) and Zonula Occludens-1 (ZO1), as well as the desmosomal protein Desmoglein-2 (DSG2). CLDN3 forms transmembrane sealing strands that anchor to the actin cytoskeleton via ZO1 [18, 26] whereas DSG2 provides mechanical strength through desmosomal adhesion [23, 24]. In the most distal colon (Figure 3a-f), all three markers are decreased in samples from untreated *Hol^Tg/Tg^*mice compared to WT, consistent with the massive bacterial infiltration detected in this region (Figure 2c,d). Prior GDNF treatment of *Hol^Tg/Tg^* mice between P4-P8 increases immunofluorescence signal intensity for all three markers, reaching intermediate levels between untreated mutants and WT controls for CLDN3 and ZO1, and slightly exceeding WT levels for DSG2 (Figure 3a-f). In the proximal colon (Figures 3g-i and S2), CLDN3 and ZO1 levels are again reduced in untreated *Hol^Tg/Tg^* mice relative to WT but not as much as in the distal colon, in agreement with the more modest bacterial infiltration detected in proximal colon (Figure 2e). DSG2 behaves differently in this segment, being now slightly increased in untreated *Hol^Tg/Tg^* mice relative to WT. Yet, all three markers are again increased after GDNF treatment, reaching almost WT levels for CLDN3 (not significant) and ZO1, or again slightly exceeding WT levels for DSG2 (Figures 3g-i and S2).

**Figure 3.**
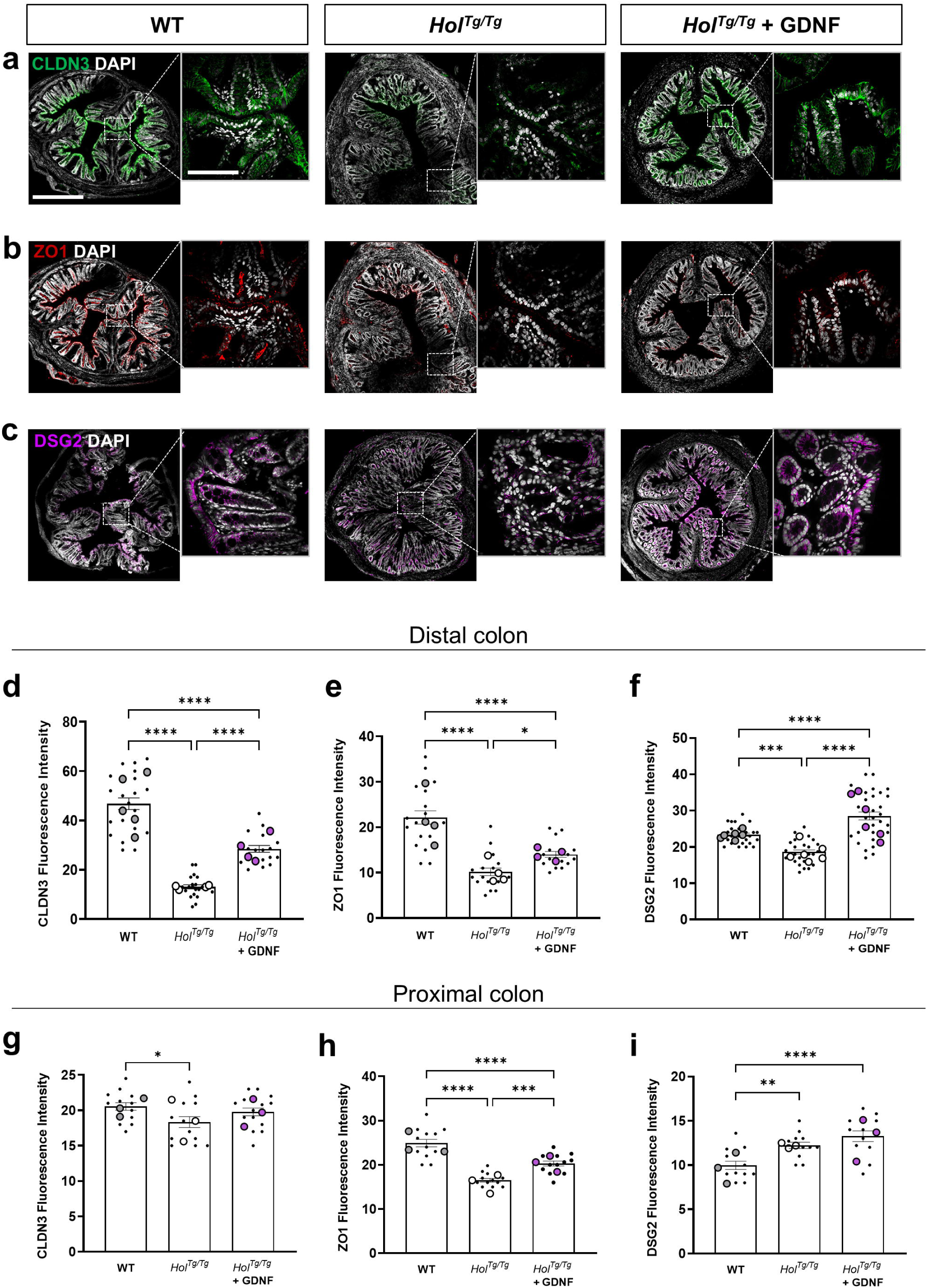
GDNF treatment enhances the expression of key junction proteins in the colonic epithelium of P20 *Hol^Tg/Tg^* mice. (**a-f**) Representative immunofluorescence staining images (a-c) and accompanying quantitative analyses (d-f) showing that GDNF treatment partially restores protein levels of epithelial CLDN3 (a,d), ZO1 (b,e) and DSG2 (c,f) in the distal colon of *Hol^Tg/Tg^*mice. (**g-i**) Quantitative analysis showing a similar outcome in proximal colon samples, for which representative images are displayed in Fig.S2. Representative images in panels a-c correspond to 15µm-thick z-stack projections (scale bar, 300 µm). Fluorescence intensity in panels d-i is expressed in arbitrary units, with large dots indicating the mean value for each mouse (n=4-5 mice per group)) and small black dots indicating each microscopic field analyzed (3-5 fields of view per animal). *p < .05, **p < .01, ***p < .001, ****p < .0001; one-way ANOVA with Tukey’s post-hoc test.

As epithelial barrier disruption and ensuing bacterial translocation should trigger local inflammation in the colon, we then immunostained colon cross-sections of P20 mice to assess neutrophil infiltration in the mucosa. Neutrophils are major innate immune cells recruited early during inflammation and release myeloperoxidase [52], an enzyme that generates reactive oxygen species (ROS) to combat pathogens [53–55]. As expected, samples from *Hol^Tg/Tg^* mice exhibit an increased number of MPO+ cells compared to WT mice in both the proximal (Figure S3a,b) and distal colon (Figure S3c,d), although it is ∼5X more pronounced in the distal segment (72 ± 7 cells/mm^2^ in proximal *vs*. 390 ± 51 cells/mm^2^ in distal). Prior GDNF treatment of *Hol^Tg/Tg^* mice markedly decreases the number of MPO+ cells close to WT-like levels in both colon segments, consistent with an anti-inflammatory potential in the context of HSCR (Figure S3), as previously reported in IBD models [27, 29]. Altogether, these results demonstrate that rectal GDNF treatment between P4-P8 improves epithelial barrier integrity and reduces neutrophil infiltration in the colon of *Hol^Tg/Tg^* mice at P20, not only at the primary site of the disease in the distal segment but also in the proximal segment with overtly normal ENS density.

### GDNF treatment has a broad effect on innate and adaptive immune responses in the colon of *Hol^Tg/Tg^* mice, with a predominance of the innate branch

To verify the effect of GDNF treatment on the gastrointestinal immune system in a more comprehensive manner, we next turned to a multicolor flow cytometry approach that we recently developed for profiling both the innate and adaptive branches [51]. Using 26 markers in accordance to our previously described gating strategies [51], we systematically quantified 55 lymphoid (Figures 4-7 and S4-S7; summarized in Figures 11-12 and Table 1) and 17 myeloid cell populations (Figures 4,8-10 and S8; summarized in Figures 11-12 and Table 2). Like the epithelial barrier studies described above, we analyzed both proximal and distal colon segments of P20 *Hol^Tg/Tg^* mice that were treated or not with GDNF between P4-P8, again using WT mice as reference control. We identified five distinct patterns, four being associated with the response to GDNF treatment (in efficacy order): (1) complete return to normal in both segments (*i.e.*, WT-like levels in both proximal and distal colon), (2) complete return to normal in one segment only (*i.e.*, WT-like levels in either proximal or distal colon), (3) incomplete return to normal in any segment (*i.e.*, partial return to WT levels in proximal and/or distal colon), and (4) no normalizing effect of GDNF treatment in any segment. The fifth pattern corresponds to immune cell populations that are not altered in the HSCR context, where given cell frequencies in *Hol^Tg/Tg^*mice remain stable and similar to WT controls. Of note, when an immune cell population exhibited two distinct patterns in proximal *vs*. distal segments, we prioritized the best response for categorization purposes. For example, when complete normalization occurs in only one segment (pattern 2), the other segment may show partial normalization, no effect of GDNF, or no alteration by HSCR (patterns 3–5).

**Figure 4.**
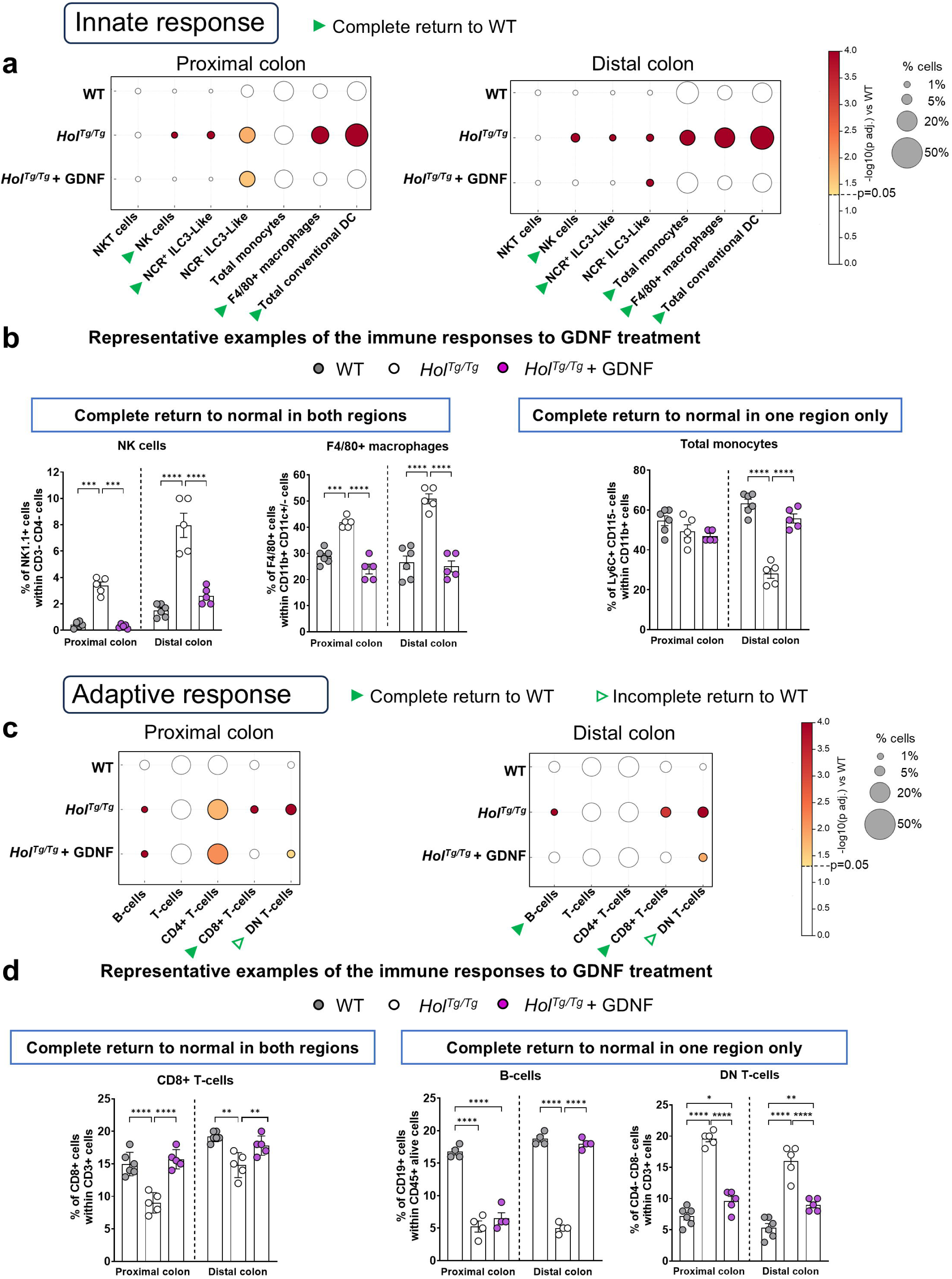
GDNF treatment influences both innate and adaptive immune responses in the entire colon of P20 *Hol^Tg/Tg^* mice. (**a,c**) Dot plots showing relative differences of indicated innate (a) and adaptive (b) immune cell populations in the proximal (left panel) and distal (right panel) colon from WT and *Hol^Tg/Tg^* mice treated or not with GDNF (n=5-6 mice per group). (**b,d**) Representative examples of the diverse responses to GDNF treatment for indicated cell populations (other graphs can be found in Fig.S4). Green arrowheads point to significant changes induced by GDNF treatment, with filled and empty arrowheads indicating either complete or partial return to WT levels, respectively. *p < .05, **p < .01, ***p < .001, ****p < .0001; two-way ANOVA with Sidak’s post-hoc test.

**Figure 5.**
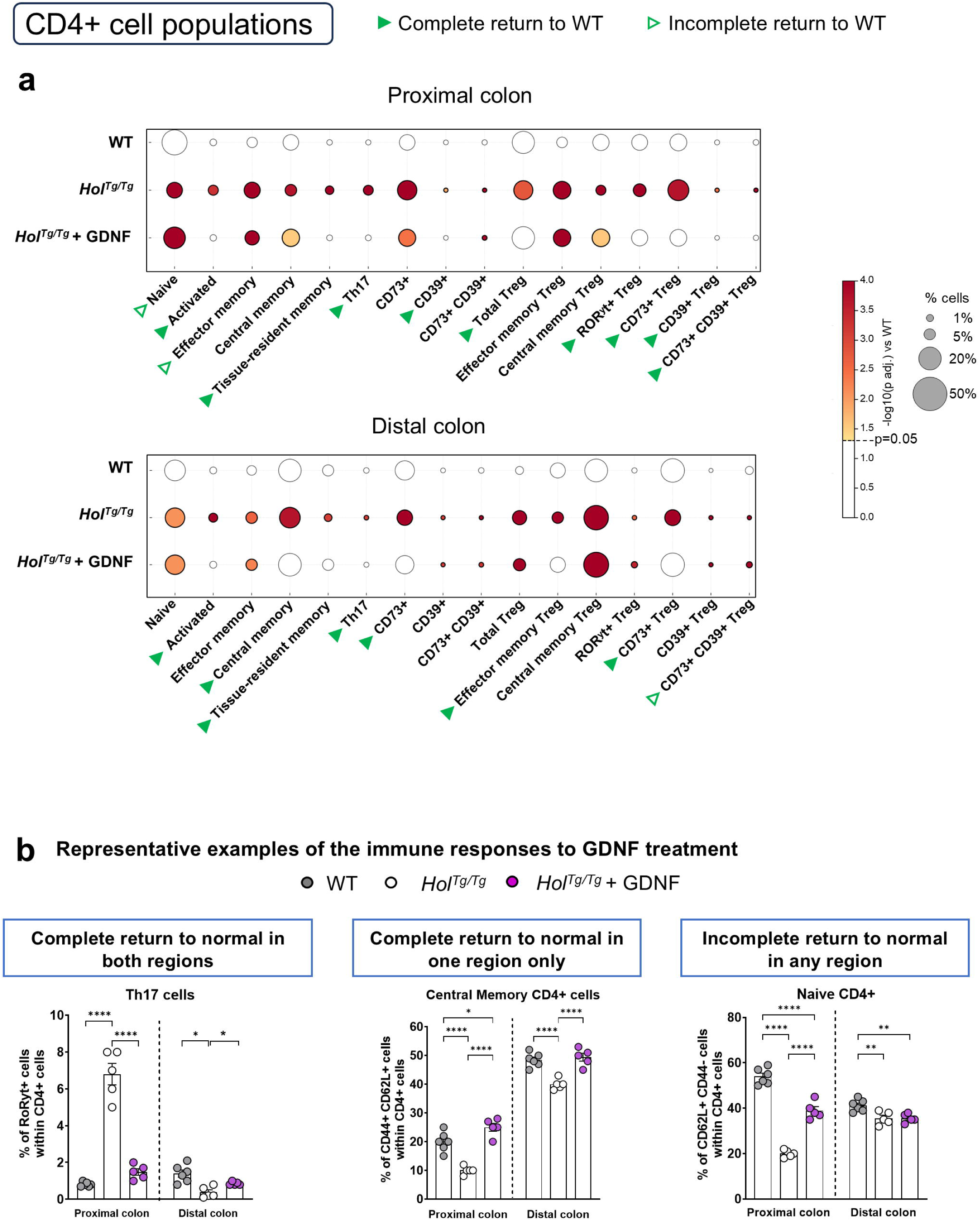
GDNF treatment impacts multiple CD4⁺ T-cell subsets in at least one colonic region of P20 *Hol^Tg/Tg^* mice. (**a**) Dot plots showing relative differences of indicated CD4⁺ T-cell subsets in the proximal (left panel) and distal (right panel) colon from WT and *Hol^Tg/Tg^* mice treated or not with GDNF (n=5-6 mice per group). (**b**) Representative examples of the diverse responses to GDNF treatment for indicated cell populations (other graphs can be found in Fig.S5). Green arrowheads point to significant changes induced by GDNF treatment, with filled and empty arrowheads indicating either complete or partial return to WT levels, respectively. ***p < .001, ****p < .0001; two-way ANOVA with Sidak’s post-hoc test.

**Figure 6.**
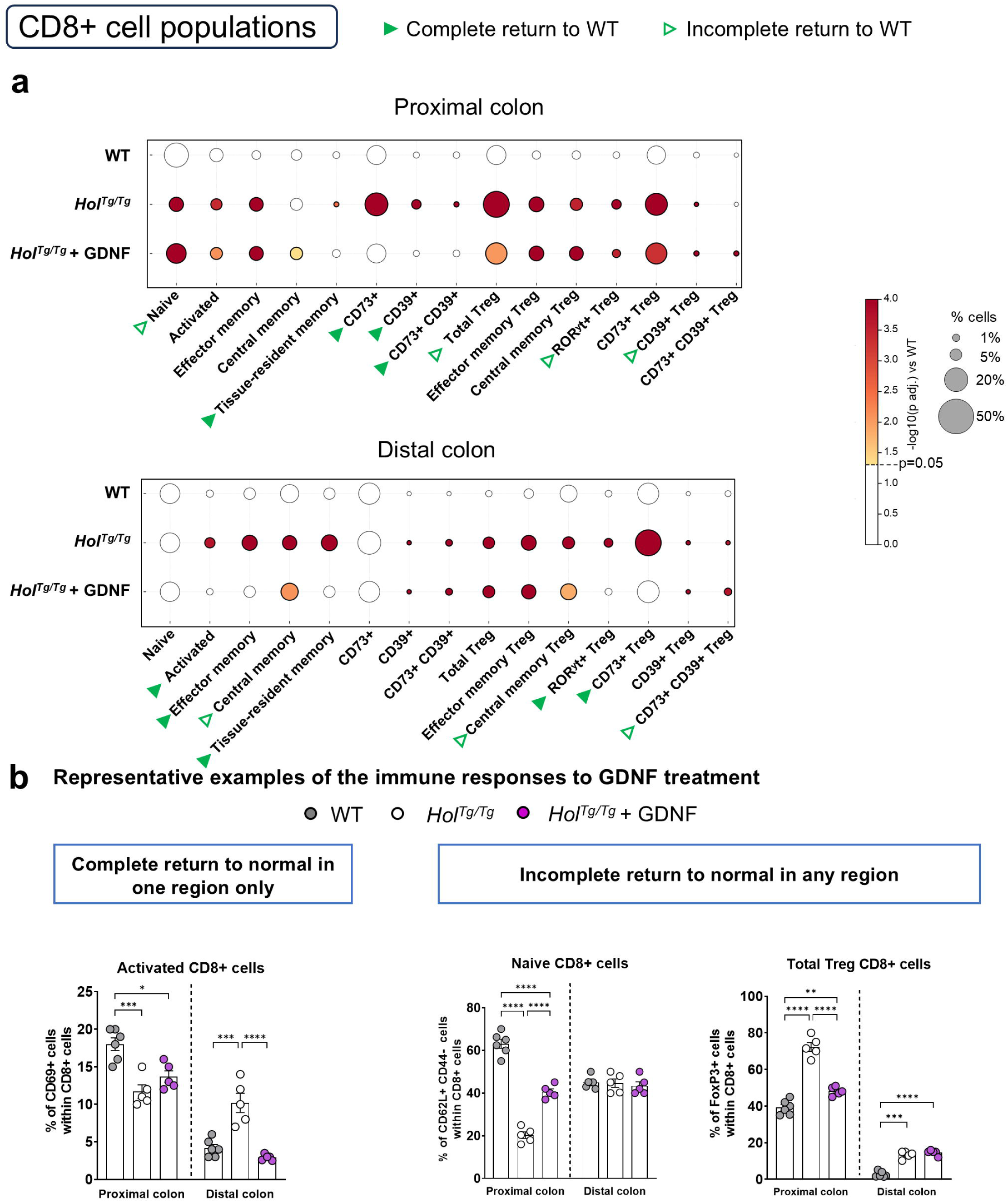
GDNF treatment impacts multiple CD8⁺ T-cell subsets in at least one colonic region of P20 *Hol^Tg/Tg^*mice. (**a**) Dot plots showing relative differences of indicated CD8⁺ T-cell subsets in the proximal (left panel) and distal (right panel) colon from WT and *Hol^Tg/Tg^* mice treated or not with GDNF (n=5-6 mice per group). (**b**) Representative examples of the diverse responses to GDNF treatment for indicated cell populations (other graphs can be found in Fig.S6). Green arrowheads point to significant changes induced by GDNF treatment, with filled and empty arrowheads indicating either complete or partial return to WT levels, respectively. **p < .01, ***p < .001, ****p < .0001; two-way ANOVA with Sidak’s post-hoc test.

**Figure 7.**
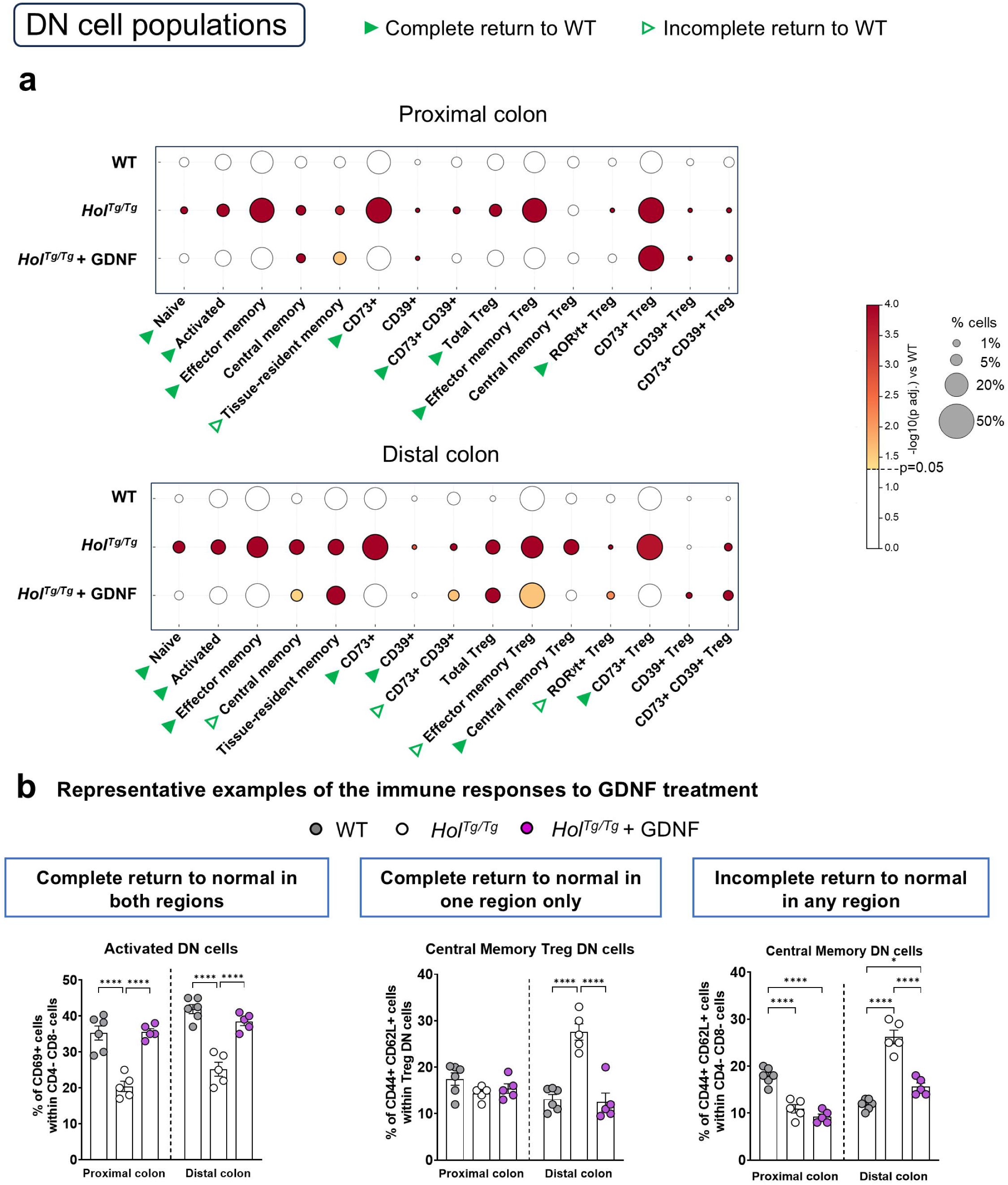
GDNF treatment impacts multiple CD4-CD8 double-negative T-cell subsets in at least one colonic region of P20 *Hol^Tg/Tg^* mice. (**a**) Dot plots showing relative differences of indicated CD4-CD8 double-negative T-cell subsets in the proximal (left panel) and distal (right panel) colon from WT and *Hol^Tg/Tg^* mice treated or not with GDNF (n=5-6 mice per group). (**b**) Representative examples of the diverse responses to GDNF treatment for indicated cell populations (other graphs can be found in Fig.S7). Green arrowheads point to significant changes induced by GDNF treatment, with filled and empty arrowheads indicating either complete or partial return to WT levels, respectively. **p < .01, ***p < .001, ****p < .0001; two-way ANOVA with Sidak’s post-hoc test.

**Figure 8.**
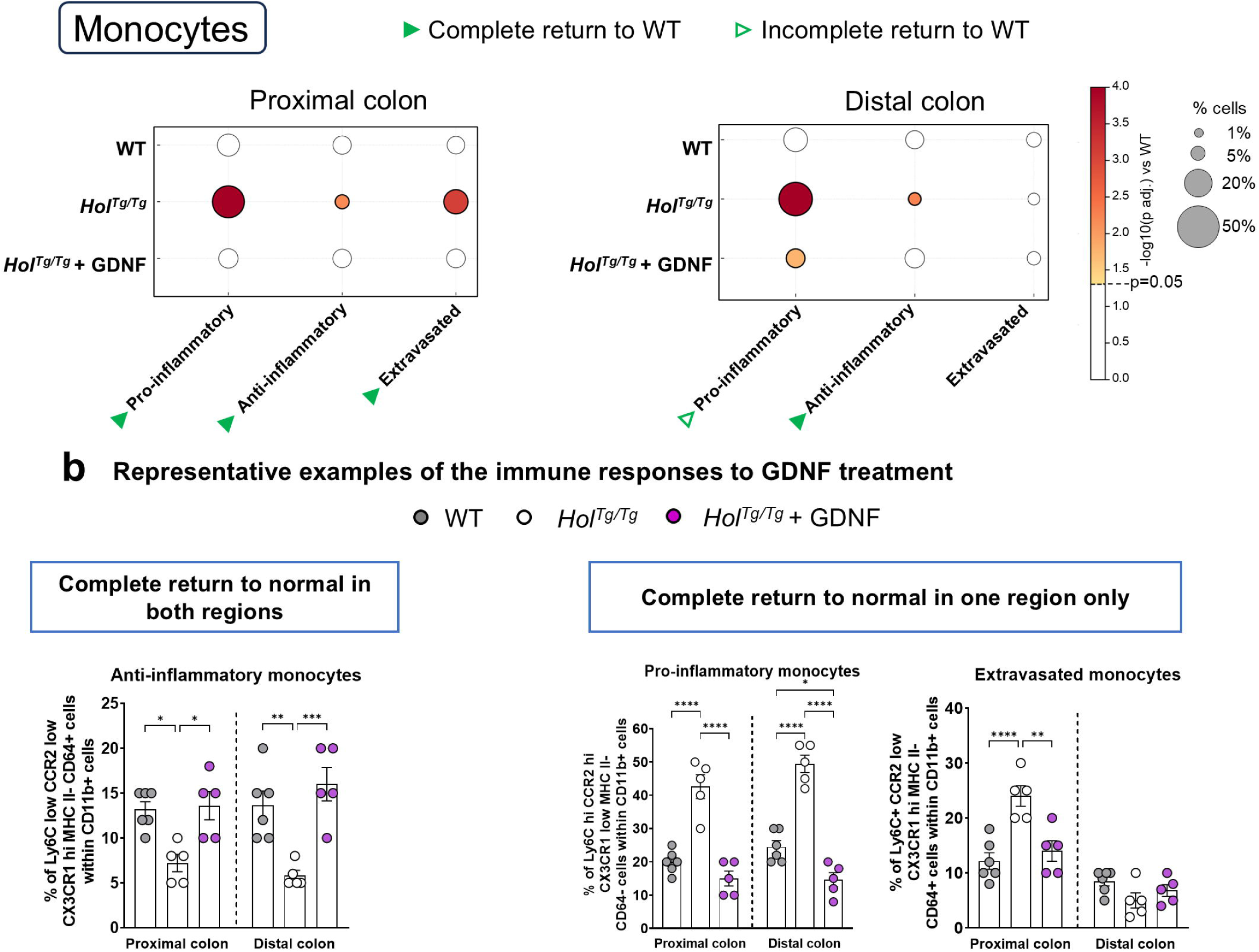
GDNF treatment impacts multiple monocyte subsets in at least one colonic region of P20 *Hol^Tg/Tg^*mice. (**a**) Dot plots showing relative differences of indicated monocyte subsets in the proximal (left panel) and distal (right panel) colon from WT and *Hol^Tg/Tg^* mice treated or not with GDNF (n=5-6 mice per group). (**b**) Representative examples of the diverse responses to GDNF treatment for indicated cell populations. Filled green arrowheads point to significant changes induced by GDNF treatment, with complete return to WT levels. *p < .05, **p < .01, ***p < .001, ****p < .0001; two-way ANOVA with Sidak’s post-hoc test.

**Figure 9.**
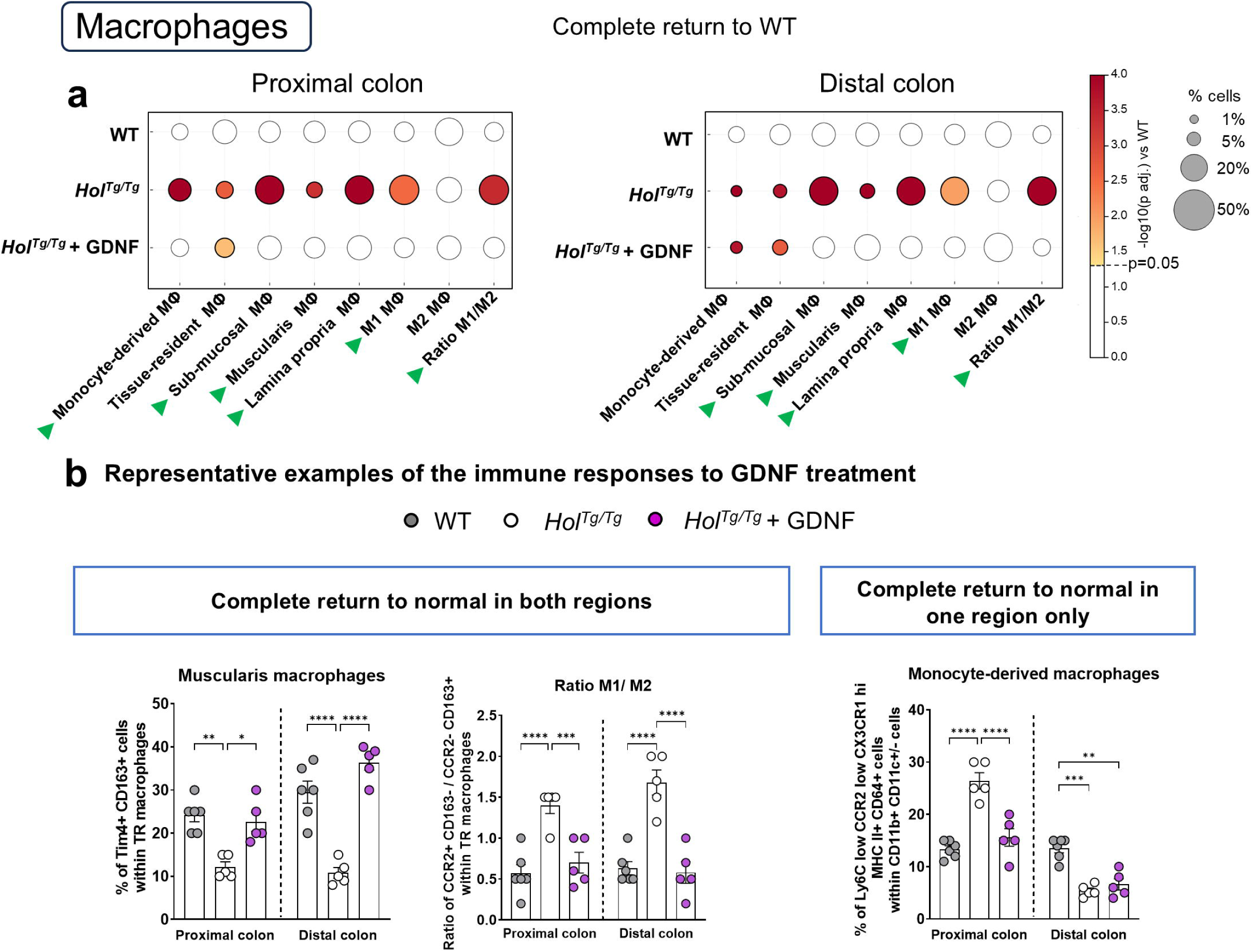
GDNF treatment impacts multiple macrophage subsets in at least one colonic region of P20 *Hol^Tg/Tg^*mice. (**a**) Dot plots showing relative differences of indicated macrophage subsets in the proximal (left panel) and distal (right panel) colon from WT and *Hol^Tg/Tg^* mice treated or not with GDNF (n=5-6 mice per group). (**b**) Representative examples of the diverse responses to GDNF treatment for indicated cell populations (other graphs can be found in Fig.S8a). Filled green arrowheads point to significant changes induced by GDNF treatment, with complete return to WT levels. *p < .05, **p < .01, ***p < .001, ****p < .0001; two-way ANOVA with Sidak’s post-hoc test.

**Figure 10.**
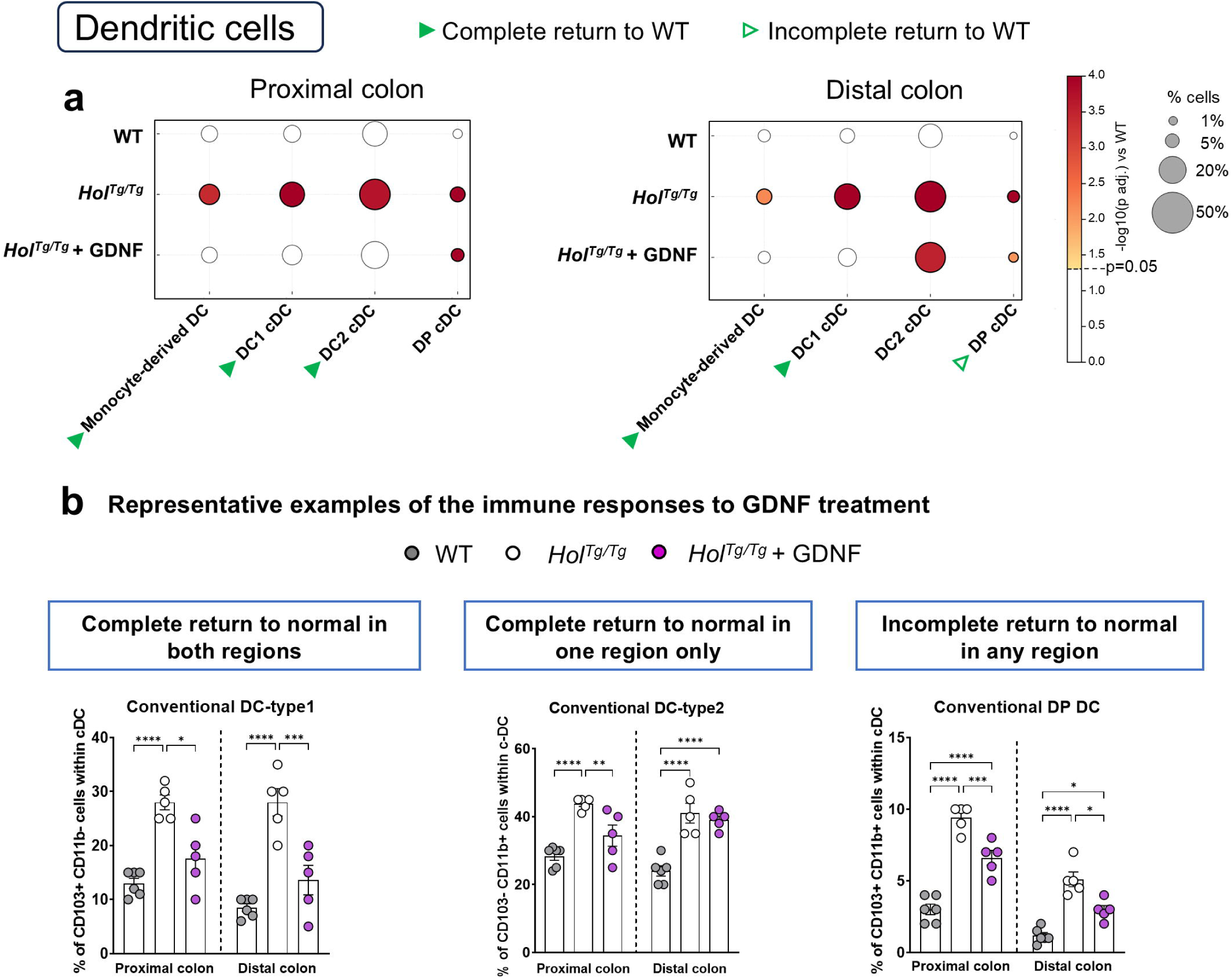
GDNF treatment impacts multiple dendritic cell subsets in at least one colonic region of P20 *Hol^Tg/Tg^* mice. (**a**) Dot plots showing relative differences of indicated dendritic cell subsets in the proximal (left panel) and distal (right panel) colon from WT and *Hol^Tg/Tg^* mice treated or not with GDNF (n=5-6 mice per group). (**b**) Representative examples of the diverse responses to GDNF treatment for indicated cell populations (other graphs can be found in Fig.S8b). Green arrowheads point to significant changes induced by GDNF treatment, with filled and empty arrowheads indicating either complete or partial return to WT levels, respectively. *p < .05, ***p < .001, ****p < .0001; two-way ANOVA with Sidak’s post-hoc test.

**Figure 11.**
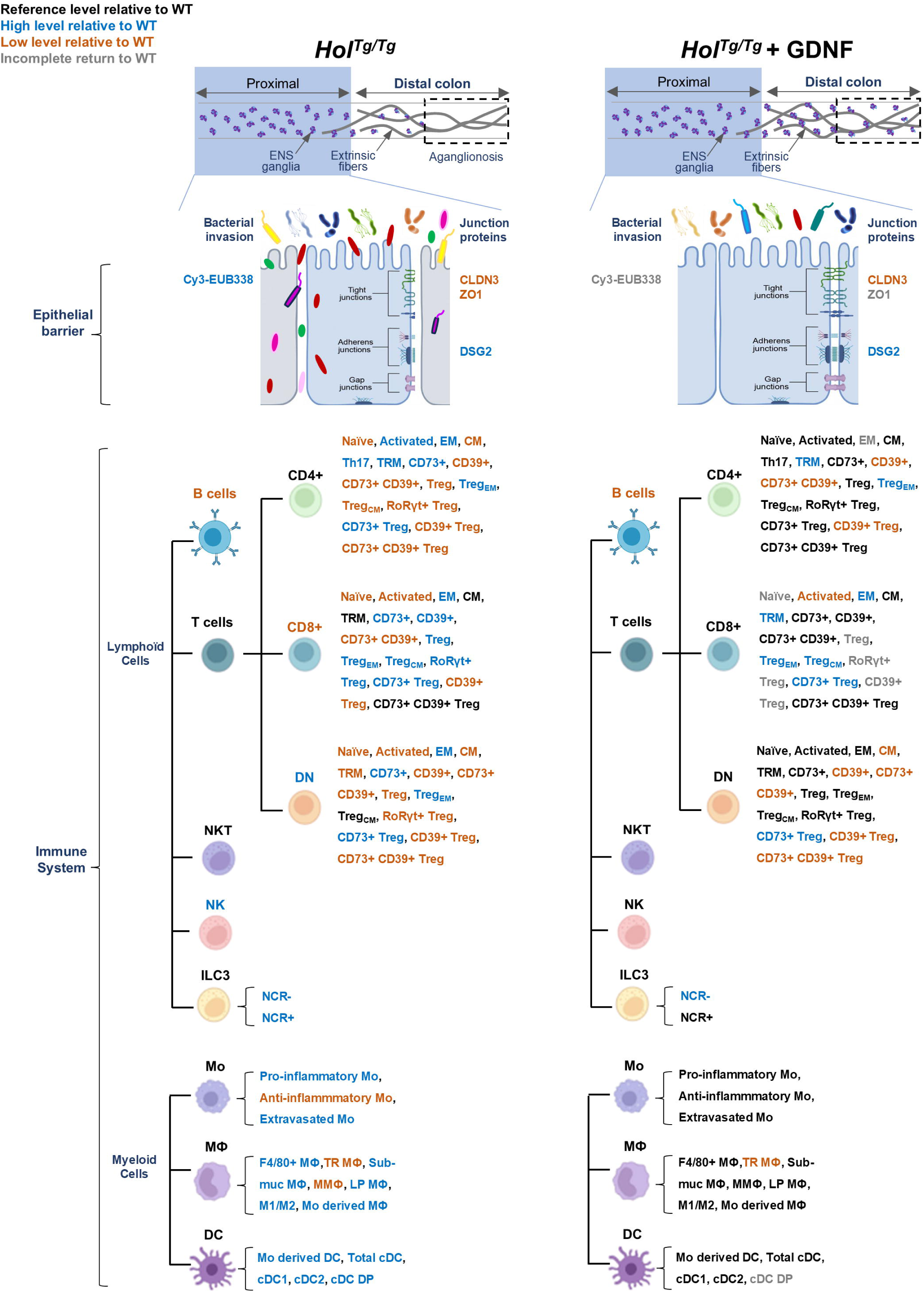
Overview of GDNF treatment effects on epithelial and immune cells from the normo-ganglionic proximal colon of P20 *Hol^Tg/Tg^* mice. For all markers or cell subsets, black text indicates normal WT-like levels, while blue and orange text indicate higher or lower expression than WT, respectively. Grey text indicates an incomplete return toward WT-like levels after GDNF treatment.

**Figure 12.**
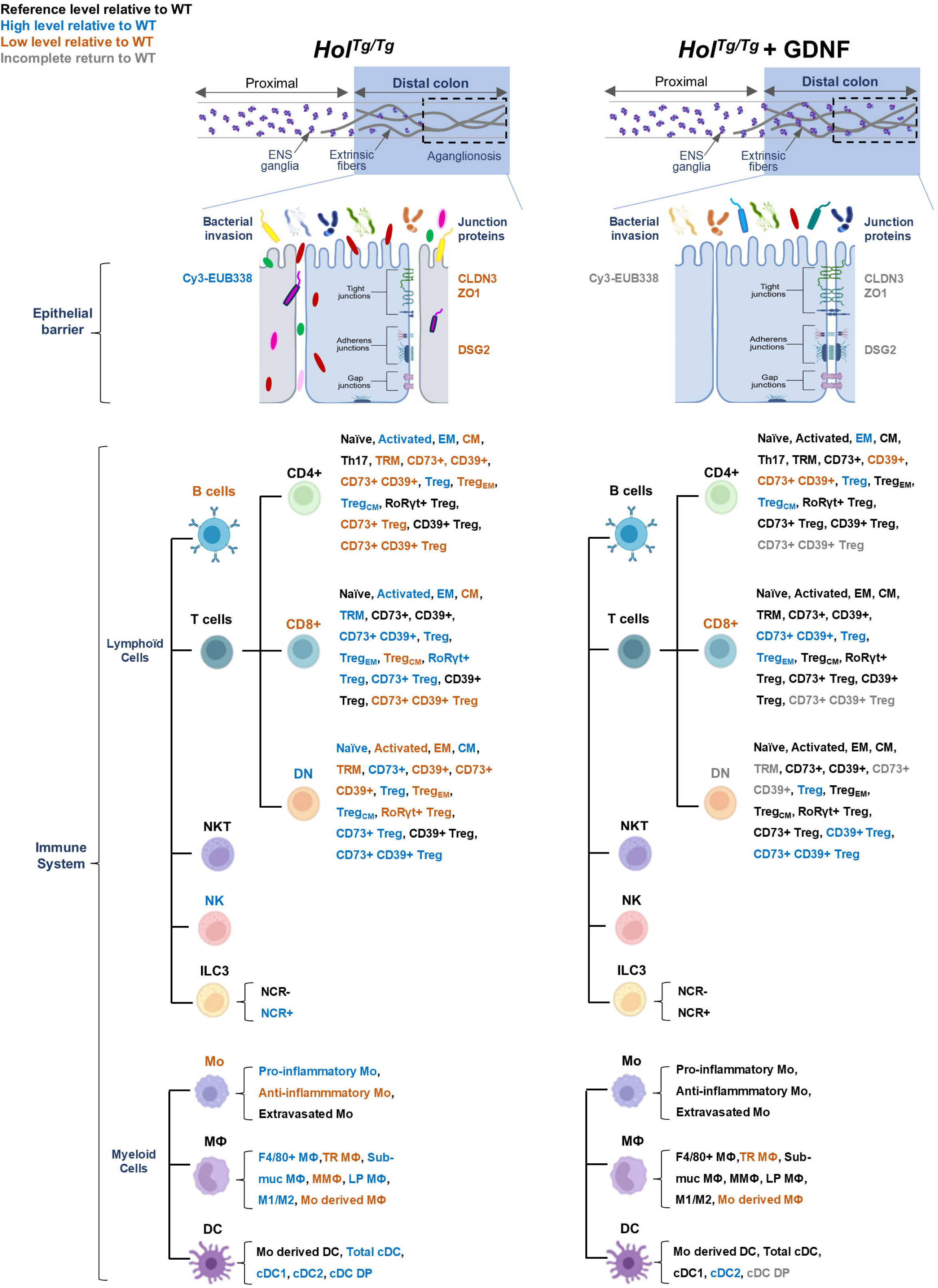
Overview of GDNF treatment effects on epithelial and immune cells from the hypo/aganglionic distal colon of P20 *Hol^Tg/Tg^* mice. For all markers or cell subsets, black text indicates normal WT-like levels, while blue and orange text indicate higher or lower expression than WT, respectively. Grey text indicates an incomplete return toward WT-like levels after GDNF treatment.

**Table 1:**
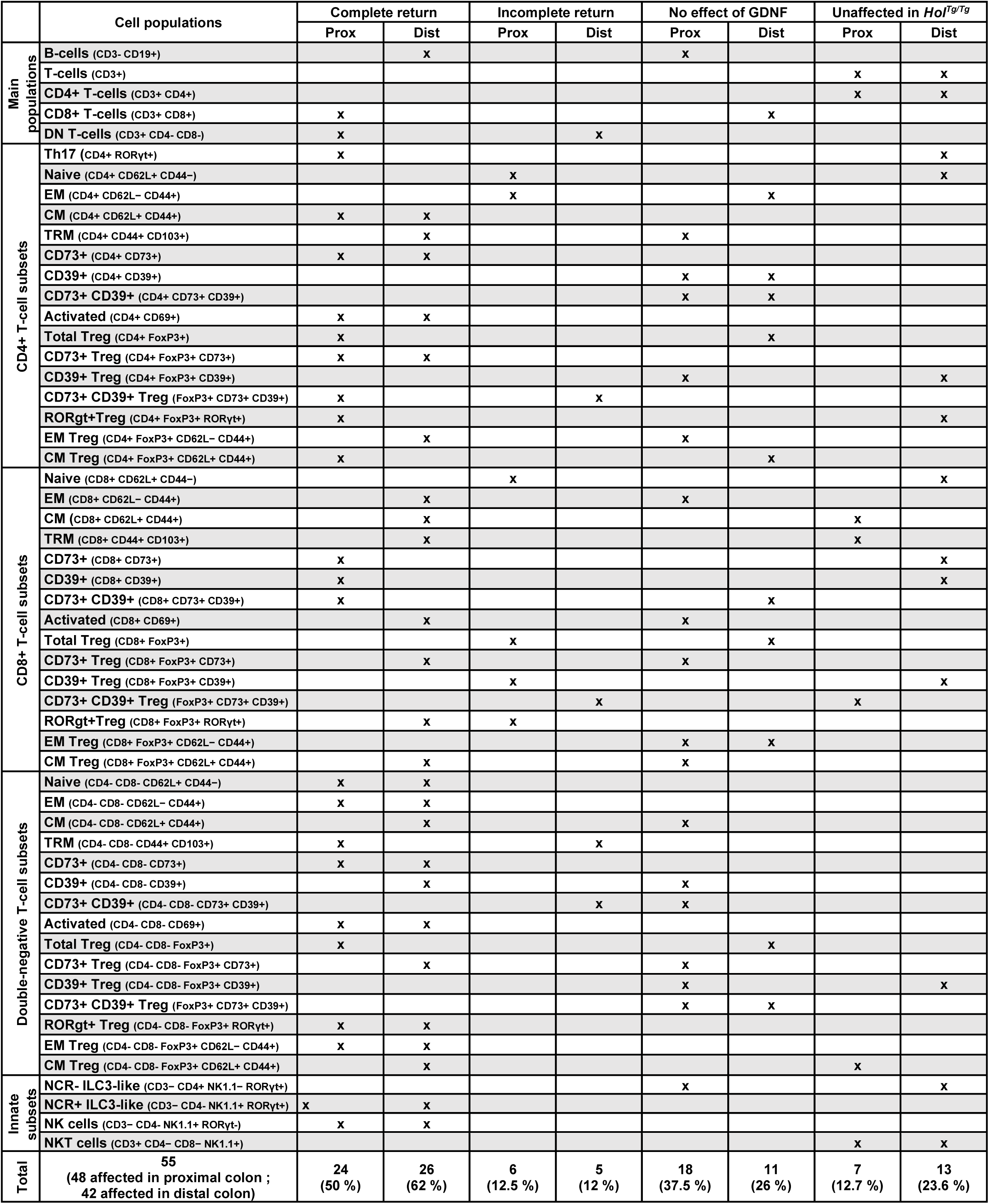
Effect of GDNF on altered lymphoid populations.

**Table 2:**
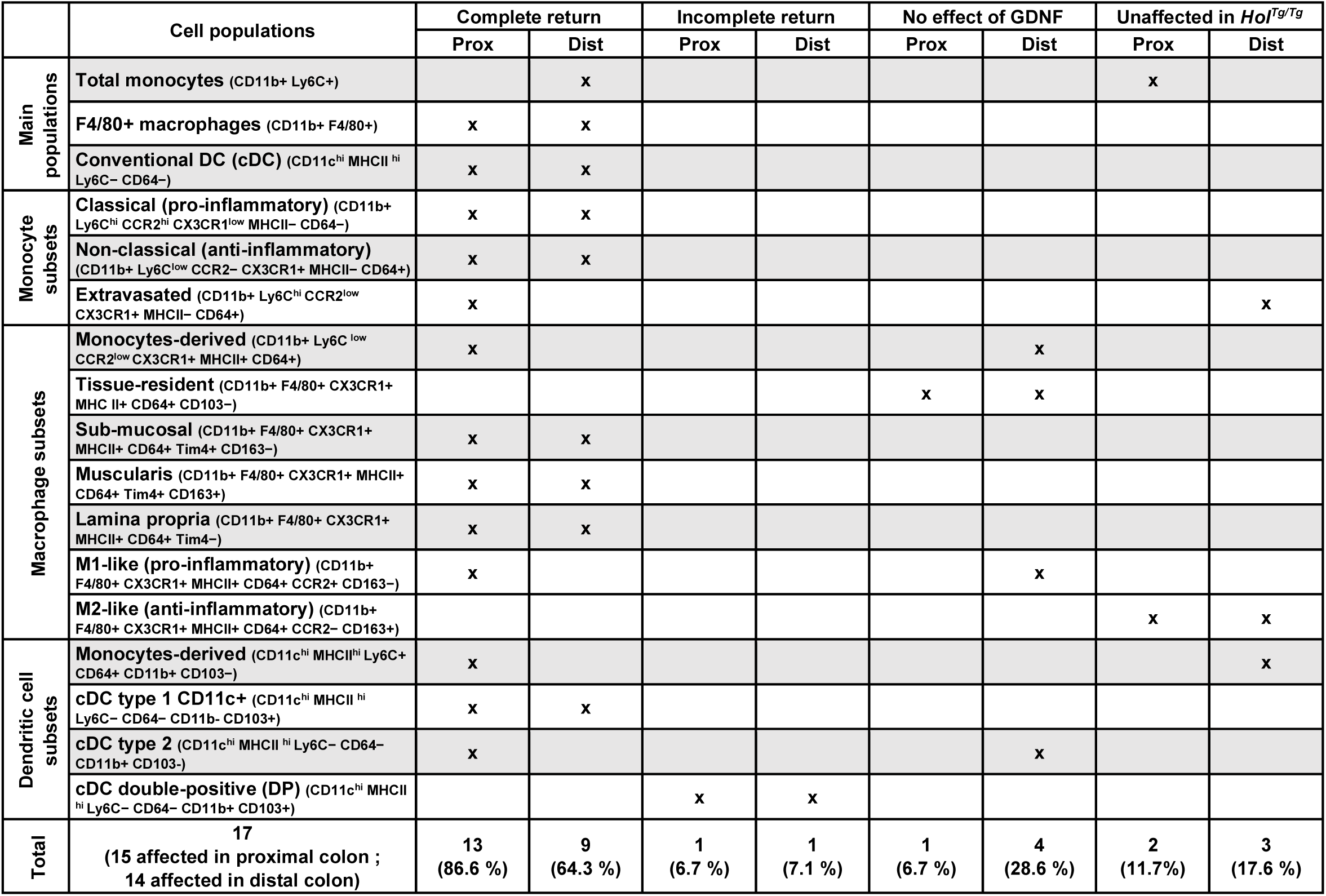
Effect of GDNF on altered myeloid populations.

To provide rapid non-specific defense, the innate immune compartment uses a large variety of cell types that include natural killer (NK) cells (CD3^-^ CD4^-^ NK1.1^+^ RORgt^-^), natural killer T-cells (NKT) (CD3^+^ CD4^-^ CD8^-^ NK1.1^+^), innate lymphoid cells type 3 (ILC3) expressing (CD3^-^ CD4^-^ NK1.1^+^ RORgt^+^) or not (CD3^-^ CD4^+^ NK1.1^-^ RORgt^+^) the natural cytotoxicity receptor (NCR), and myeloid cells such as monocytes (CD11b^+^ Ly6C^+^), macrophages (CD11b^+^ F4/80^+^), and dendritic cells (CD11c^Hi^ MHCII^Hi^) [56]. In comparison, the adaptive immune system responds more slowly, but its B (CD19^+^) and T (CD3^+^) lymphocytes provide long-lasting and antigen-specific protection. When only considering these main cell populations mentioned above, we first noted that both colon segments of untreated *Hol^Tg/Tg^*mice are similarly affected compared to WT controls, with 71% (5/7) of profiled cell types affected for the innate branch and 60% (3/5) for the adaptive branch (Figures 4 and S4; summarized in Figures 11-12 and Tables 1-2). Out of the five affected innate cell populations, four are commonly increased in both colon segments of *Hol^Tg/Tg^* mice (NK cells, NCR^+^ ILC3-like cells, total F4/80^+^ macrophages, total conventional dendritic cells) and these are accompanied by either an increase in NCR^-^ ILC3-like cells in the proximal segment or a decrease in total monocytes in the distal segment (Figures 4a,b and S4a; summarized in Figures 11-12 and Tables 1-2). All three affected adaptive cell populations are the same in both segments, two exhibiting decreased frequencies in *Hol^Tg/Tg^* mice (B-cells and CD8^+^ T-cells) while the other is increased (CD4 CD8 double-negative T-cells) (Figures 4c,d and S4b; summarized in Figures 11-12 and Table 1). In response to prior GDNF treatment, all affected innate cell populations returned to WT-like levels except NCR^-^ ILC3-like cells in the proximal segment (Figures 4a,b and S4a; summarized in Figures 11-12 and Tables 1-2). The response to GDNF treatment is less uniform for the affected adaptive cell populations, with only double-negative T-cells being corrected in both segments (but not equally), while CD8^+^ T-cells (in proximal) and B-cells (in distal) returned to WT-like levels in one segment only (Figures 4c,d and S4b; summarized in Figures 11-12 and Table 1). These results indicate that GDNF treatment can restore WT-like frequencies of several main immune cell populations, in both the innate and adaptive compartments of *Hol^Tg/Tg^* mice, with an apparently greater impact on the innate immune compartment.

### GDNF treatment differentially modulates T-cell subsets in the colon of *Hol^Tg/Tg^* mice

Applying our full panel of lymphoid cell markers [51] to all three main populations of T-cells (CD4^+^, CD8^+^, and double-negative) then revealed significant changes for several subsets that could not be anticipated when considering general markers only. This is especially true for CD4^+^ cells, which do not appear significantly affected overall in *Hol^Tg/Tg^* mice (Figures 4c,d and S4b) but which ultimately have the largest number of specific subsets affected in the context of the disease. Moreover, for both CD4^+^ and CD8^+^ cells, the number of specific subsets impacted by the disease is greater in the proximal segment (Figures 5-6 and S5-S6; summarized in Figures 11-12 and Table 1), whereas such numbers are equally high in both the proximal and distal segments for double-negative cells (Figures 7 and S7; summarized in Figures 11-12 and Table 1). As described in greater detail below for each major T-cell population, GDNF treatment nonetheless proved equally effective in restoring the frequency of more than half of the affected specific subsets to WT-like levels in both colon segments of *Hol^Tg/Tg^* mice (Figures 5-7 and S5-S7; summarized in Figures 11-12 and Table 1).

For CD4⁺ T-cells, traditionally known as helper cells, we analyzed 9 subsets of conventional naïve (CD62L^-^ CD44^-^), activated (CD69^+^), effector memory (CD62L^-^ CD44^+^), central memory (CD62L^+^ CD44^+^), tissue-resident memory (CD44^+^ CD103^+^), metabolically specialized CD73⁺ and CD39⁺ cells, as well as T helper 17 (Th17; RORgt^+^) cells. We also similarly analyzed 7 subsets of Foxp3^+^ regulatory T-cells (Treg). This analysis first revealed distinct immune profiles that markedly differ between the proximal and distal colon of untreated *Hol^Tg/Tg^*mice (Figures 5 and S5; summarized in Figures 11-12 and Table 1). In the proximal colon (left panel of Figure 5a; summarized in Figure 11 and Table 1), the observed profile is consistent with a pro-inflammatory status, characterized by increased activation of CD4⁺ T-cells, higher frequencies of Th17 cells, and a reduction in total Treg populations compared to WT mice. This is accompanied by a shift away from conventional naïve and central memory cells toward effector memory and tissue-resident memory subsets, suggesting ongoing immune stimulation. However, we also noted an enrichment in conventional CD73⁺ cells that could evoke an attempt to counterbalance inflammation. In the distal colon (right panel of Figure 5a; summarized in Figure 11 and Table 1), although CD4⁺ T-cell activation is also similarly increased, it is here accompanied by a more regulated profile, with a trend toward reduced Th17 cells (although not statistically significant) and an increase in total Treg populations compared to WT mice. Moreover, the shift from conventional central memory toward effector memory cells here appears to be much lower in amplitude compared to what we detected in the proximal segment. The distal segment also exhibits reduced frequencies of conventional tissue-resident memory cells. Intriguingly, the frequency of CD73⁺ populations changes in the opposite direction compared to the proximal segment, being reduced across both conventional and Treg compartments. Prior GDNF treatment of *Hol^Tg/Tg^* mice lead to a substantial restoration of CD4⁺ T-cell frequencies toward WT-like levels, correcting 11 out of the 16 affected subsets in the proximal segment and 7 out of 12 in the distal segment (summarized in Figures 11-12 and Table 1). Among these, four populations are fully restored in both proximal and distal colon (conventional activated, central memory and CD73^+^ cells, as well as CD73^+^ Treg) (Figures 5 and S5a). Seven subsets are fully normalized in one region only, five in the proximal colon (Th17, total Treg, central memory Treg, RORγT^+^ Treg, CD73^+^ CD39^+^ Treg) and two in the distal colon (conventional tissue-resident memory, effector memory Treg) (Figures 5 and S5b). Three showed incomplete recovery to WT-like levels in one segment, two in the proximal colon (conventional naïve and effector memory cells) and one in the distal colon (CD73^+^ CD39^+^ Treg) (Figures 5 and S5c).

For CD8⁺ T-cells, primarily known for their cytotoxic functions, we analyzed all the same subsets except the CD4-specific Th17. Interestingly, this detailed analysis of CD8⁺ subsets revealed spatially distinct immune responses that globally oppose those of CD4⁺ subsets in the colon of untreated *Hol^Tg/Tg^*mice *vs*. WT (Figures 6 and S6; summarized in Figures 11-12 and Table 1). In the proximal colon (left panel of Figure 6a; summarized in Figure 11 and Table 1), the CD8⁺ compartment appears overall more regulated in comparison to the pro-inflammatory signature of CD4⁺ subsets. Notably, we noted a reduction in activated CD8⁺ T-cells alongside a marked increase in total CD8⁺ Treg populations, including RORγt^+^ Tregs. The presence of immunosuppressive mechanisms is also supported by the enrichment in both central and effector memory Treg subsets and the increase in CD73⁺ cells in both conventional and Treg populations. In contrast, in the distal colon (right panel of Figure 6a; summarized in Figure 12 and Table 1), CD8⁺ subsets seem to be more engaged than CD4^+^ subsets in T-cell mediated inflammation. While the frequencies of activated (increased), effector memory (increased) and central memory (decreased) subsets are similarly affected for both conventional CD4^+^ and CD8^+^ cells, tissue-resident memory subsets are affected in the opposite direction, being markedly increased in the CD8^+^ compartment. However, this increased frequency of tissue-resident memory CD8^+^ cells appears nonetheless well counterbalanced by increased frequencies of multiple Treg subsets of CD8^+^ cells that are otherwise decreased or stable in corresponding CD4^+^ Treg populations (effector memory, RORγt^+^, and CD73^+^). GDNF treatment has again strong but now strictly region-specific immunomodulatory effects (Figures 6 and S6; summarized in Figures 11-12 and Table 1), as none of the CD8^+^ populations are simultaneously normalized in both proximal and distal colon of *Hol^Tg/Tg^*mice. In the proximal segment (left panel of Figure 6a), GDNF treatment restores 7 of the 12 altered subsets, including three that are fully normalized to WT-like levels (conventional CD73^+^ and/or CD39^+^) and four that exhibit partial recovery (naïve cells, total Treg, RORγt⁺ Treg, and CD39^+^ Treg). In the distal colon (right panel of Figure 6a), where 11 populations of CD8^+^ cells are affected by the disease context, GDNF treatment rescues 8, of which 7 are completely normalized (activated, effector memory, central memory, tissue-resident memory subsets of conventional CD8^+^ cells, as well as central memory, RORγt⁺ and CD73⁺ subsets of CD8^+^ Treg) and one shows incomplete recovery toward WT-like levels (CD73^+^ CD39^+^ Treg).

For CD4-CD8 double-negative T-cells, which may contribute to both cytotoxic and regulatory responses [57–59], we analyzed the same subsets as those studied for CD8^+^ cells. Like for CD4^+^ and CD8^+^ compartments, this detailed analysis of double-negative T-cells revealed several segment-specific differences in untreated *Hol^Tg/Tg^* mice (Figures 7-S7; summarized in Figures 11-12 and Table 1). However, none of the segment-specific profiles can in this case be associated with a pro-inflammatory signature. In contrast, we noted a homogeneous increase in CD73⁺ cell frequencies across both conventional and Treg subsets in both the proximal and distal colon, pointing to shared immunosuppressive mechanisms in the entire colon. In the proximal colon (left panel of Figure 7; summarized in Figure 11 and Table 1), the overall profile in *Hol^Tg/Tg^* mice appears at first glance to be similar to that of CD4^+^ T-cells, in particular for the Treg subsets. However, the conventional double-negative subsets are clearly distinguished by a reduced frequency of both activated and tissue-resident memory subsets in the context of the disease. Another notable difference is the higher frequencies of CD39^+^ cells in double-negative T-cells in general (in both conventional and Treg subsets), which are globally reduced in the HSCR context. In the distal colon (left panel of Figure 7; summarized in Figure 12 and Table 1), the overall profile of double-negative T-cells does not extensively overlap with the profile of either CD4^+^ or CD8^+^ populations for this segment. However, we noted that all three main T-cell compartments have a particularly strong enrichment in total Treg, suggesting a coordinated regulatory response. As observed for the CD4^+^ compartment, prior GDNF treatment of *Hol^Tg/Tg^* mice leads to widespread normalization of several double-negative T-cell subsets toward WT-like levels, including 6 that are fully restored in both proximal and distal colon (conventional naïve, activated, effector memory, and CD73^+^, as well as effector memory Treg and RORγt^+^ Treg). Two other subsets are fully corrected in the proximal colon only (conventional tissue-resident memory, and total Treg) whereas four others are specifically corrected in the distal segment (conventional central memory and CD39^+^, as well as central memory Treg and CD73^+^ Treg). Partial restoration is seen for only two populations, specifically in the distal colon (conventional CD73^+^ CD39^+^, and tissue-resident memory) Figure 7, Figure S7b). Hence, overall, GDNF treatment restores 8 out of 14 affected CD4-CD8 double-negative T-cell populations in the proximal colon and 12 out of 14 populations in the distal colon.

Together, these data reveal that GDNF treatment exerts broad, yet region-specific, immunomodulatory effects on all three major T-cell compartments of *Hol^Tg/Tg^* mouse colon. Out of the 55 lymphoid cell populations that were examined, 48 and 42 subsets were found to be affected by the HSCR context in the proximal and distal colon, respectively (Figures 11-12 and Table 1). In the proximal colon, GDNF treatment completely restores 50% (24/48) of the affected populations, partially corrects 12.5% (6/48), and shows no effect on 37.5% (18/48). In the distal colon, GDNF treatment leads to complete normalization for 62% (26/42) of the affected subsets, incomplete recovery for 12% (5/42), and has no effect on 26% (11/42).

### GDNF treatment extensively modulates the myeloid cell landscape in the colon of *Hol^Tg/Tg^* mice

Finally, consistent with the strong involvement of the innate response suggested upon gross analysis of main immune cell populations (Figures 4a,b and S4a), our full panel of myeloid cell markers [51] revealed robust engagement of monocyte, macrophage, and dendritic cell subsets in the colon of *Hol^Tg/Tg^* mice and good responsiveness to GDNF treatment. In contrast to lymphoid cell subsets, the observed profiles here appear more homogeneous in both the proximal and distal colon, with untreated *Hol^Tg/Tg^* mice generally exhibiting a pro-inflammatory profile that shifts toward an anti-inflammatory profile after GDNF treatment (Figures 8-10 and S8; summarized in Figures 11-12 and Table 2).

For monocytes, which can give rise to both macrophages and dendritic cells, we analyzed three subsets known as extravasated (CD11b^+^ Ly6C^+(hi)^ CCR2^+(lo)^ CX3CR1^+^ MHCII^-^ CD64^+^), classical/pro-inflammatory (CD11b^+^ Ly6C^+(hi)^ CCR2^+(hi)^ CX3CR1^+(lo)^ MHCII^-^ CD64^-^), and non-classical/anti-inflammatory (CD11b^+^ Ly6C^+(lo)^ CCR2^-^ CX3CR1^+^ MHCII^-^ CD64^+^) (Figure 8). In both colon segments of untreated *Hol^Tg/Tg^* mice, we found that the frequencies of classical/pro-inflammatory and non-classical/anti-inflammatory monocytes are increased and decreased, respectively, reflecting a global inflammatory state. Yet, extravasated monocytes recruited from the circulation are selectively increased in the proximal colon. This localized accumulation supports the notion that a particularly pronounced inflammatory process is ongoing in the proximal segment, as also observed for CD4⁺ T-cell subsets (Figures 5 and S5). Importantly, all of these affected monocyte subsets are normalized back to WT-like levels after GDNF treatment (Figure 8).

For macrophages, the professional phagocytes, we analyzed 7 subsets known as monocyte-derived (CD11b^+^ Ly6C^+(lo)^ CCR2^+(lo)^ CX3CR1^+^ MHCII^+^ CD64^+^), tissue-resident (CD11b^+^ F4/80^+^ CX3CR1^+^ MHCII^+^ CD64^+^ CD103^-^), lamina propria (CD11b^+^ F4/80^+^ CX3CR1^+^ MHCII^+^ CD64^+^ Tim4^-^), submucosal (CD11b^+^ F4/80^+^ CX3CR1^+^ MHCII^+^ CD64^+^ Tim4^+^ CD163^-^), muscularis (CD11b^+^ F4/80^+^ CX3CR1^+^ MHCII^+^ CD64^+^ Tim4^+^ CD163^+^), pro-inflammatory M1 (CD11b^+^ F4/80^+^ CX3CR1^+^ MHCII^+^ CD64^+^ CCR2^+^ CD163^-^) and anti-inflammatory M2 (CD11b^+^ F4/80^+^ CX3CR1^+^ MHCII^+^ CD64^+^ CCR2^-^ CD163^+^) (Figures 9 and S8). In both proximal and distal colon of untreated *Hol^Tg/Tg^* mice, the frequencies of lamina propria and submucosal macrophages are increased, while those of tissue-resident and muscularis macrophages are significantly diminished. The M1/M2 ratio also appears similarly elevated in both colon segments (1.4 ± 0.22 in proximal; 1.68 ± 0.34 in distal), mostly due to the marked expansion of the M1 pool. Only one regional difference is noted, again in the sense of more pronounced inflammation in the proximal colon, where the frequencies of monocyte-derived macrophages are increased while they are in contrast reduced in the distal segment. Most of these changes noted above are fully corrected after GDNF treatment (Figures 9 and S8). *Hol^Tg/Tg^* mice that were previously administered GDNF enemas have WT-like levels for M1/M2 ratios and frequencies of lamina propria, submucosal and muscularis macrophages in both colon segments, whereas the frequency of monocyte-derived macrophages is fully rescued in proximal colon only.

For dendritic cells, the accessory cells specialized in antigen presentation, we analyzed four subsets known as monocyte-derived (CD11c^+(hi)^ MHCII^+(hi)^ Ly6C^+^ CD64^+^ CD11b^+^ CD103^-^), conventional type 1 (CD11c^+(hi)^ MHCII^+(hi)^ Ly6C^-^ CD64^-^ CD11b^-^ CD103^+^), conventional type 2 (CD11c^+(hi)^ MHCII^+(hi)^ Ly6C^-^ CD64^-^ CD11b^+^ CD103^-^), and conventional double-positive (CD11c^+(hi)^ MHCII^+(hi)^ Ly6C^-^ CD64^-^ CD11b^+^ CD103^+^) (Figures 10 and S8). The frequencies of all three conventional dendritic cell subsets are increased in untreated *Hol^Tg/Tg^* mice across both colon regions (Figure 10, Figure S8c). As observed for monocyte-derived macrophages (Figure 9), the frequency of monocyte-derived dendritic cells is increased selectively in the proximal colon (Figure 10) – a finding also in line with the increased frequencies of extravasated monocytes in this colon segment only (Figure 8). GDNF treatment corrects the frequencies of conventional type 1 (fully) and conventional double-positive (partially) subsets toward WT-like levels in both colon segments, whereas those of monocyte-derived and conventional type 2 subsets are corrected in the proximal colon only.

Altogether, these results demonstrate that GDNF treatment can broadly mitigate the pro-inflammatory myeloid landscape associated with the HSCR context. Out of 17 myeloid populations that were analyzed in total,15 and 14 are affected by the disease in the proximal and distal colon, respectively (Figures 11-12 and Table 2). In the proximal colon, prior GDNF treatment of *Hol^Tg/Tg^*mice completely restores 86.6% (13/15) of these affected populations, partially corrects 6.7% (1/15), and has no effect on 6.7% (1/15). In the distal colon, GDNF treatment leads to complete normalization for 64.3% (9/14) of affected subsets, incomplete recovery for 7.1% (1/14), and shows no effect on 28.6% (4/14).

## Discussion

The main goal of this study was to better understand the therapeutic effects of GDNF enemas for treating HSCR. Building on previous work showing that GDNF treatment can promote enteric neurogenesis and improve colonic motility in various HSCR mouse models [10, 11, 15], our new findings provide additional insights into its broader restorative effects on epithelial and immune cells that also likely contribute to the increased lifespan of GDNF-treated HSCR mice. In doing so, we provide here the most comprehensive analysis to date of the immune system from the colon of a HSCR mouse model. As discussed in greater detail below, another major outcome of the current study is the demonstration that GDNF treatment positively impacts these additional non-ENS cell populations not only at the primary site of the disease in the distal colon but also in the proximal colon. Our data thus add to the notion that the entire gastrointestinal ecosystem is perturbed in the ENS-containing regions upstream of the aganglionic segment, in both HSCR mice [10, 42, 60–70] and human patients [42, 62, 71–73], further suggesting that GDNF-based therapy could decrease most if not all complications attributed to global dysfunction of these upstream regions after standard HSCR surgery.

### GDNF treatment of *Hol^Tg/Tg^* mice flattens region-specific differences in colonic permeability: is the epithelial stem cell niche involved?

Our epithelial barrier data from untreated *Hol^Tg/Tg^*mice revealed that junctional proteins are most severely impaired in the aganglionic distal segment, consistent with the massive bacterial translocation and neutrophil infiltration also observed in this region. Although less severely affected, the proximal colon displayed similar alterations of all three parameters. These findings in mice are in line with two recent human studies that analyzed different sets of junctional proteins in the colon of children with HSCR [71], suggesting widespread alterations of epithelial junctions in both hypo/aganglionic and normo-ganglionic regions. The only protein analyzed in all three studies is CLDN3, which however was found not to be altered in the two human studies. This discrepancy could be explained either by species-specific differences or, more likely, by the experimental setting. Notably, the smaller size of mouse samples allows for a complete circumferential analysis of the colonic mucosa, which is not possible with larger human samples. Since our data indicate that tight junction abnormalities are not uniformly distributed around the circumference, this means that these circumferencial differences might go unnoticed when analyzing small quadrants of human colon cross-sections.

Most interestingly, we found that GDNF treatment of *Hol^Tg/Tg^*mice reduces bacterial infiltration in both the proximal and distal colon to the same extent, regardless of the differing severity otherwise observed in corresponding segments of untreated animals. As the associated restoration of junctional proteins in mature epithelial cells is observed 12 days (at P20) after the end of the GDNF treatment (between P4-P8), such protective effect against bacterial translocation cannot be explained by an action of GDNF on these mature epithelial cells – in which only the enterochromaffin lineage is under direct GDNF/RET regulation [32, 33]. Instead, we think it is more plausible that GDNF treatment promotes global improvement of epithelial regeneration, a key process in epithelial homeostasis that has recently been reported to be impaired in the colon of humans and mice with HAEC [62]. Indeed, mouse colonic epithelial cells undergo complete turnover every 5–7 days [74, 75], meaning that the epithelium has been entirely renewed almost twice since the treatment window ended. Given that the impaired epithelial regeneration in HAEC has been associated with reduced numbers of LGR5^+^ stem cells [62], it is particularly interesting to note that systemic administration of GDNF was recently shown to promote mucosal healing by increasing *Lgr5* gene expression and stem cell proliferation in the context of DSS-induced ulcerative colitis [28]. These observations strengthen the notion that HAEC is much closer to IBD than initially thought, suggesting that new insights gained from one of these conditions could clarify our understanding of the other, including for treatment options.

Importantly, GDNF-induced repair of the intestinal stem cell niche and delayed impact on the mature epithelial cells by 5-7 days help to explain our previous observations that His-tagged GDNF continuously accumulated in the distal colon wall during the P4-P8 treatment, while extending this window until P12 had no additional benefit on *Hol^Tg/Tg^* survival [10]. Moreover, as the enema volume we used can reach the proximal ENS-containing colon of *Hol^Tg/Tg^* pups [10], the barrier defect newly detected in this region helps to understand other previous observations in GDNF-treated *Hol^Tg/Tg^*animals. Indeed, direct entry of GDNF in the proximal colon could underly the correction of the cholinergic *vs*. nitrergic neuron imbalance in this region as well as the widespread autoregulation of endogenous GDNF-RET that was detected across the entire length of the colon [10]. This autoregulation capacity of the GDNF-RET pathway, also described in other contexts [76, 77], is particularly important when evaluating the therapeutic value of GDNF enemas for the treatment of HSCR. Indeed, RET signaling is often reduced but not abrogated in affected children [78, 79], suggesting that this positive feedback loop could sufficiently enhance RET signaling in such cases where initial RET levels are low. Although RET is dispensable for GDNF-induced neurogenesis, which we found instead occurs via NCAM1 signaling [15], current knowledge suggests that GDNF-induced mucosal healing is RET-dependent. Whether GDNF acts directly or indirectly is however less clear. Blocking RET signaling with the chemical inhibitor BLU-667 was recently shown to impair GDNF-induced activation of the stem cell niche in differentiated human organoids [28]. Yet, in mice, although the *Ret* gene has been reported to be highly transcribed in the stem cell niche [31], detectable levels of mechanotransduction-mediated RET protein phosphorylation have been observed in only a small subset of activated stem cells [35]. Converging evidence suggests that GDNF-RET signaling could perhaps more globally stimulate intestinal stem cells indirectly, by activating WNT signaling in the niche [28, 34]. Targeting this pathway makes sense in the treatment of HSCR, as stromal cells that release WNT signals have been implicated in the pathological cascade leading to HAEC [62]. Moreover, long lasting effects could be provided by the newly GDNF-induced and self-sustaining ENS [10, 15], which is known to influence many aspects of intestinal stem cell proliferation and differentiation via both neuronal [80, 81] and glial [82, 83] pathways. Not to mention the positive impact of GDNF treatment on the microbiota [10], which is also known to influence intestinal stem cell biology during early postnatal development [84]. More work will definitely be needed to disentangle all these possibilities.

### GDNF treatment has widespread anti-inflammatory effects in the entire colon of *Hol^Tg/Tg^* mice

Based on the more severe epithelial barrier defects observed in the distal colon of untreated *Hol^Tg/Tg^* mice, we initially anticipated that this hypo/aganglionic segment would exhibit more pronounced pro-inflammatory immune activation than in the proximal normo-ganglionic colon. However, our detailed flow cytometry analyses revealed the opposite, in line with previous studies reporting more severe HAEC in proximal than distal colon [42]. Indeed, for both innate and adaptive responses, we found that untreated *Hol^Tg/Tg^*mice show more numerous shifts in immune cell frequencies and stronger signatures of ongoing inflammation in the proximal colon. In the distal colon, our data rather suggest engagement of regulatory mechanisms to tolerate and limit excessive inflammation at P20, as notably highlighted by the general increase in total Treg frequencies for all three major subpopulations of T-cells (CD4^+^, CD8^+^, CD4-CD8 double-negative). The less leaky proximal colon appears to be better able to naturally tolerate persistent inflammatory processes at the same P20 stage. Regardless of the colon segment, we further noted that innate immune cell subsets are generally more profoundly affected than adaptive populations in untreated *Hol^Tg/Tg^* mice. Given the hierarchical nature of immune responses, this robust engagement of innate immunity might have led, at least in part, to the observed changes of adaptive immune cell frequencies in these mice.

Several important specific findings previously reported by others using fewer markers in either *Ednrb*-mutant mice or human patient samples were replicated in the current comprehensive analysis of untreated *Hol^Tg/Tg^*mice, suggesting that these observations might be generalizable for HSCR/HAEC. In mice, this includes a global decrease in B-cells [85] and a global increase in pro-inflammatory M1 macrophages [42]. This latter observation was also noted in human patients [42], while the study of another cohort [43] reported the same aganglionic colon-specific increase in total Treg mentioned above for *Hol^Tg/Tg^*mice. Interestingly, increased frequencies of M1 macrophages [86] and Treg [87] are also hallmarks of IBD. Consistent with the pathological overlap between both conditions, other important changes in untreated *Hol^Tg/Tg^* mice not previously reported by others in the context of HSCR/HAEC nonetheless replicated previous findings made in the context of IBD. This includes increased frequencies of dendritic cells [88, 89] as well as increased expression of CD73 [90], which mirrors the overall increased frequencies of CD73+ T-cells that we observed in *Hol^Tg/Tg^* mice

In accordance with a study in IBD mice [91], we found that GDNF treatment of *Hol^Tg/Tg^* mice reduces the frequency of pro-inflammatory M1 macrophages and thereby normalizes M1/M2 ratios in both proximal and distal colon. Our comprehensive analysis revealed that many other pro-inflammatory immune cell subsets are also similarly decreased after GDNF treatment, including classical/pro-inflammatory monocytes, antimicrobial NCR^+^ ILC3-like cells and Th17 cells (in proximal colon only; not engaged in distal colon). In addition, GDNF treatment of *Hol^Tg/Tg^*mice globally increases the frequency of non-classical/anti-inflammatory monocytes, but not for M2 macrophages – which do not significantly vary in any experimental group. Another particularly striking effect of GDNF treatment is the nearly global restoration of CD73⁺ T-cell populations. However, given that CD73, an ectonucleotidase responsible for adenosine production, is a key immunosuppression regulator [92, 93], this observation could only reflect a healthier environment. Determining whether GDNF-induced changes in immune cell subset frequencies are an integral part of the treatment or simply a consequence of it is complex. Time-resolved profiling studies, both during and after GDNF treatment, will be needed to get a clearer picture. Determining whether GDNF acts directly on immune cells or indirectly via the newly induced ENS will be equally challenging. As for epithelial cells, the newly GDNF-induced ENS could very well influence immune cell frequencies and/or functions via cytokines and neurotransmitters released by either neurons [94–97] or glia [98, 99]. Yet, it is worth remembering that RET is present in a large array of immune cell populations [44] and that GDNF is administered when the mucosal immune system is immature [100, 101]. This suggests that early postnatal treatment with GDNF between P4-P8 might accelerate the maturation of some RET^+^ immune cells involved in balancing reactivity and tolerance at later stages. This would explain why the immune profile of GDNF-treated *Hol^Tg/Tg^* mice is overall generally anti-inflammatory despite the presence of invading bacteria that are not completely blocked after treatment. Furthermore, here again, a potential indirect contribution from the microbiota of GDNF-treated *Hol^Tg/Tg^* mice should not be neglected, especially when considering the major role played by the microbiota in shaping immune tolerance [102, 103].

## Conclusion

This study reinforces GDNF’s status as a promising treatment for HSCR and potentially other gastrointestinal disorders with overlapping pathophysiology. We showed that GDNF treatment has pleiotropic benefits extending well beyond its neurogenic action, in both the proximal and distal colon. This strongly suggests that GDNF treatment of short-segment HSCR would not only prevent the need for surgery in the distal aganglionic colon but also reduce long-term problems attributed to proximal colon dysfunction. However, we also note that GDNF treatment is not fully effective at preventing bacterial translocation and re-establishing immune homeostasis. This means that other factors affecting these parameters could help to develop a more effective combination therapy. Diet-derived factors seem a promising avenue in this regard, as food can have a profound influence on mucosal biology and survival of HSCR mice despite aganglionosis [60].

## Limitations of the study

To reduce the number of animals used in our research, we did not include a PBS enema control group. Yet, we know from our prior work that this control group is indistinguishable from untreated *Hol^Tg/Tg^* mice in terms of animal survival [10]. Moreover, we used data on mucosal bacterial density as an indicator of epithelial barrier permeability, but we did not measure it directly in the proximal colon using a functional test, as we had done previously for the distal colon [10]. Quantification of systemic inflammation (*e.g*., in blood) would also help assess the broader, systemic impact of GDNF treatment rather than only its localized intestinal effects. Finally, the whole study has been performed using a single mouse model, and thus these data will need to be replicated in other models and/or considered in the context of future human clinical trials.

## Data availability statement

The data that support the findings of this study are available in the Materials and Methods, Results, and/or Supplemental Material of this article.

## Author contributions

Conceived the study: NP and RS. Supervised the study: NP, RS and MAJ. Designed the experiments: NL, AY and NP. Performed the experiments: NL (all, except for Figures 3 and S2), JT and ZG (Figures 3 and S2), AG (some of the treatments in mice). Analyzed the data: NL, RS, MAJ and NP. Contributed reagents/materials/analysis tools: NP and MAJ. Wrote the paper: NL and MAL (first draft), RS (preliminary review and editing), and NP (final review and editing). All authors read and approved the final manuscript.

## Acknowledgements

The authors gratefully acknowledge the Cellular analyses and Imaging core (CERMO-FC, UQAM) for their assistance with confocal microscopy and flow cytometry data acquisition and analysis. This work was supported by grants from the Canadian Institutes of Health Research (CIHR # PJT-180290) and the SynergiQc program from the *Consortium québécois de découverte du medicament* (CQDM #SYN-329) to NP. MAJ holds the Tier 2 CIHR Canada Research Chair in Human Immuno-Virology. NL and AY were supported by scholarships from the CERMO-FC and the Fonds de recherche du Québec – Santé (FRQS), respectively.

