## Supplementary material for "GDNF enemas improve epithelial and immune defects in both aganglionic and ganglionic colon of Hirschsprung mice": Fig.S

Lassoued *et al*.

**SUPPLEMENTAL MATERIAL**

**Figure S1. Supplemental data showing that GDNF treatment reduces both local and peripheral bacterial translocation in P20 *Hol*$^{Tg/Tg}$ mice**

**Figure S2. Supplemental data showing that GDNF treatment enhances the expression of key junction proteins in the colonic epithelium of P20 *Hol*$^{Tg/Tg}$ mice**

**Figure S3. GDNF treatment limits the infiltration of MPO$^{+}$ cells in both proximal and distal colon of P20 *Hol*$^{Tg/Tg}$ mice**

**Figure S4. Supplemental data showing that GDNF treatment influences both innate and adaptive immune responses in the colon of P20 *Hol*$^{Tg/Tg}$ mice**

**Figure S5. Supplemental data showing that GDNF treatment impacts multiple CD4$^{+}$ T-cell subsets in at least one colonic region of P20 *Hol*$^{Tg/Tg}$ mice**

**Figure S6. Supplemental data showing that GDNF treatment impacts multiple CD8$^{+}$ T-cell subsets in at least one colonic region of P20 *Hol*$^{Tg/Tg}$ mice**

**Figure S7. Supplemental data showing that GDNF treatment impacts multiple CD4-CD8 double-negative T-cell subsets in at least one colonic region of P20 *Hol*$^{Tg/Tg}$ mice**

**Figure S8. Supplemental data showing that GDNF treatment impacts multiple macrophage and dendritic cell subsets in at least one colonic region of P20 *Hol*$^{Tg/Tg}$ mice**

**Table S1: List of antibodies and FISH probes utilized**

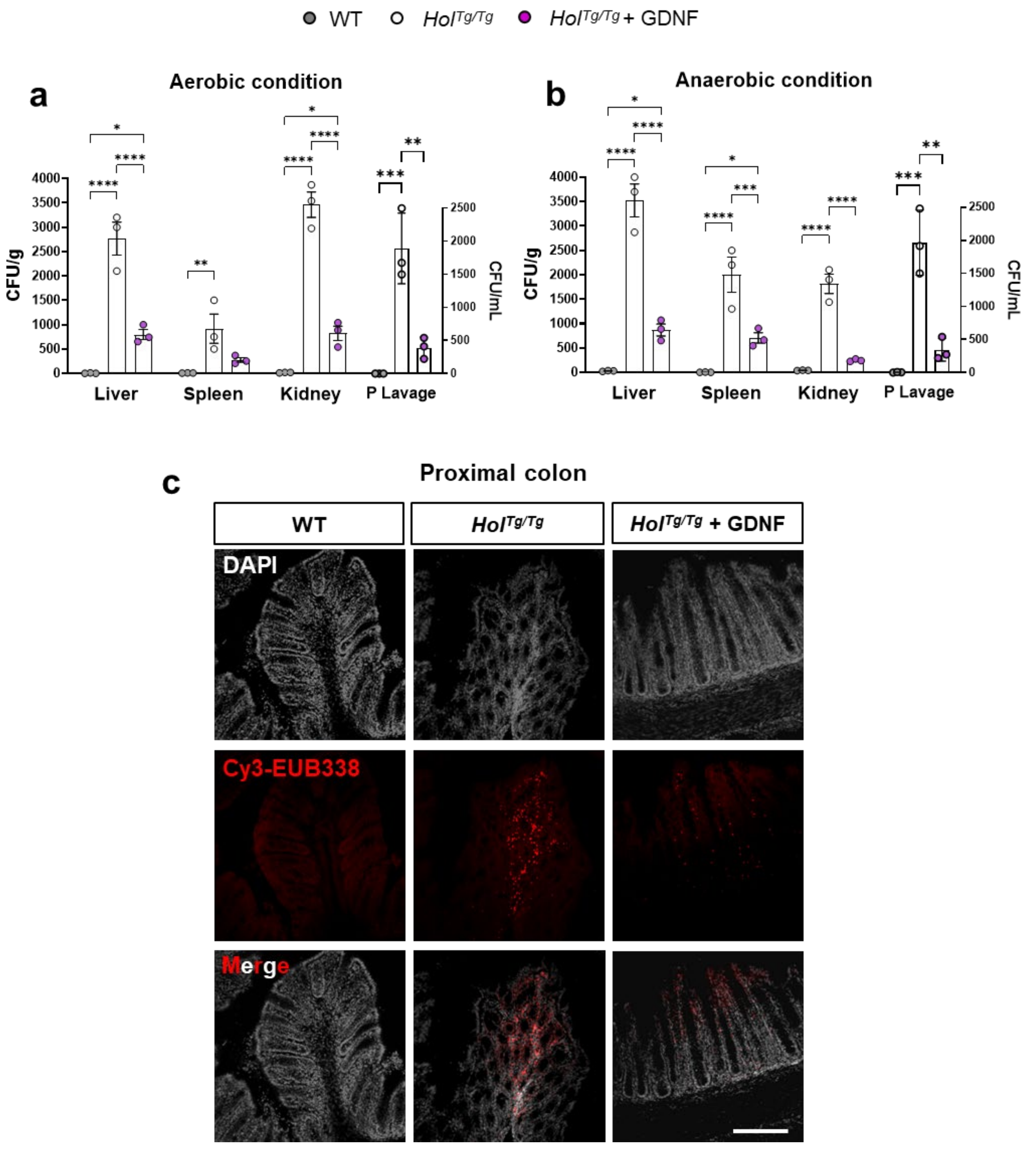


**Figure S1. Supplemental data showing that GDNF treatment reduces both local and peripheral bacterial translocation in P20 *$Hol^{Tg/Tg}$* mice.** (**a**,**b**) Quantitative analysis of bacterial load in samples of peritoneal lavage and peripheral organs (liver, spleen and kidneys) that were cultured on chocolate agar medium under aerobic (a) and anaerobic (b) conditions. GDNF treatment decreases back to WT-like levels the otherwise elevated bacterial load in samples from *$Hol^{Tg/Tg}$* mice (n=5 mice per group). (**c**) FISH staining of bacterial 16S rRNA showing that GDNF treatment limits bacterial translocation into the colonic mucosa of proximal colon from *$Hol^{Tg/Tg}$* mice. These representative images correspond to 15µm-thick z-stack projections (scale bar, 70 µm). *$p < .05$, **$p < .01$, ***$p < .001$, ****$p < .0001$; Two-way ANOVA with Tukey's post-hoc test. *In support of main Fig.2.*

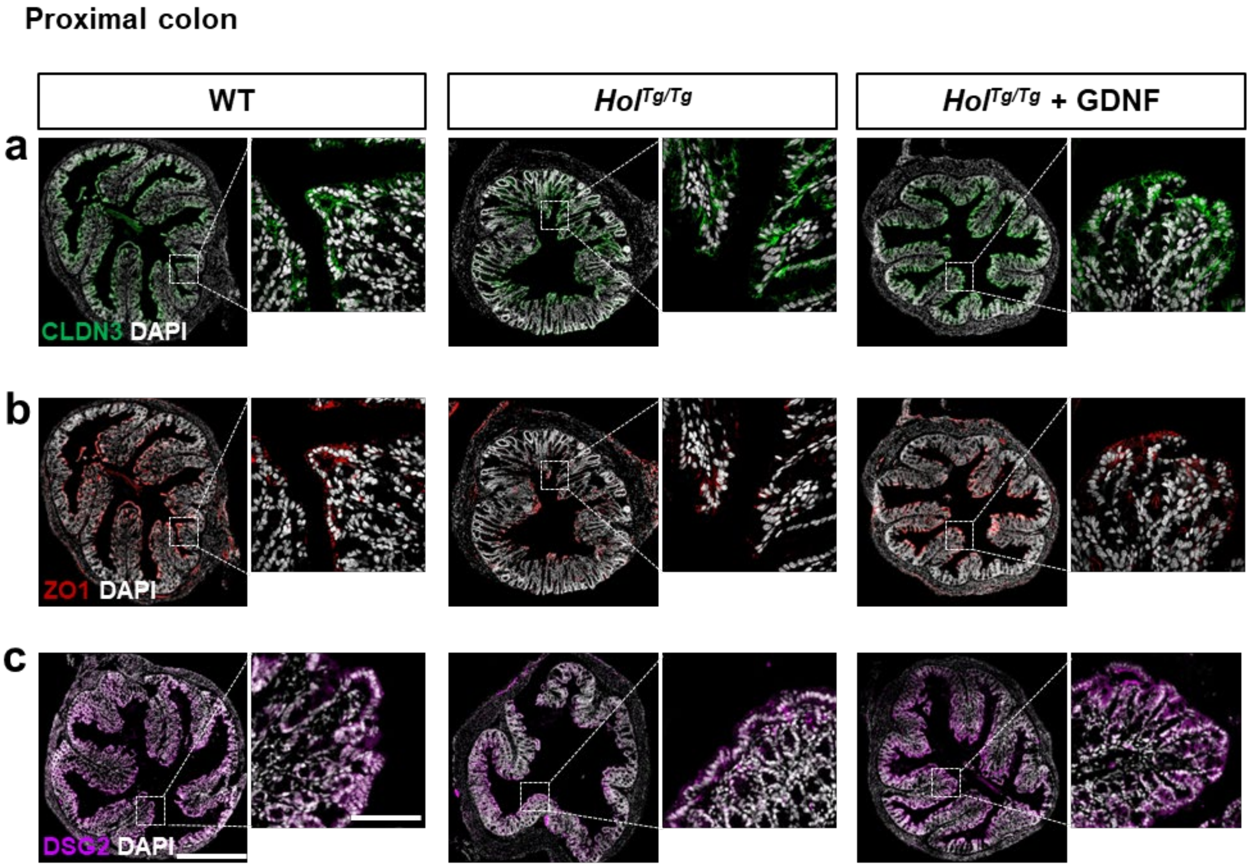


**Figure S2. Supplemental data showing that GDNF treatment enhances the expression of key junction proteins in the colonic epithelium of P20 *Hol*$^{Tg/Tg}$ mice**. (**a-c**) Representative immunofluorescence images of the epithelial junction proteins CLDN3 (a), ZO1 (b) and DSG2 (c) in normo-ganglionic proximal colon of WT and *Hol*$^{Tg/Tg}$ mice treated or not with GDNF. All images show a 15µm-thick z-stack projections. Scale bar, 300 µm. *In support of main Fig.3.*

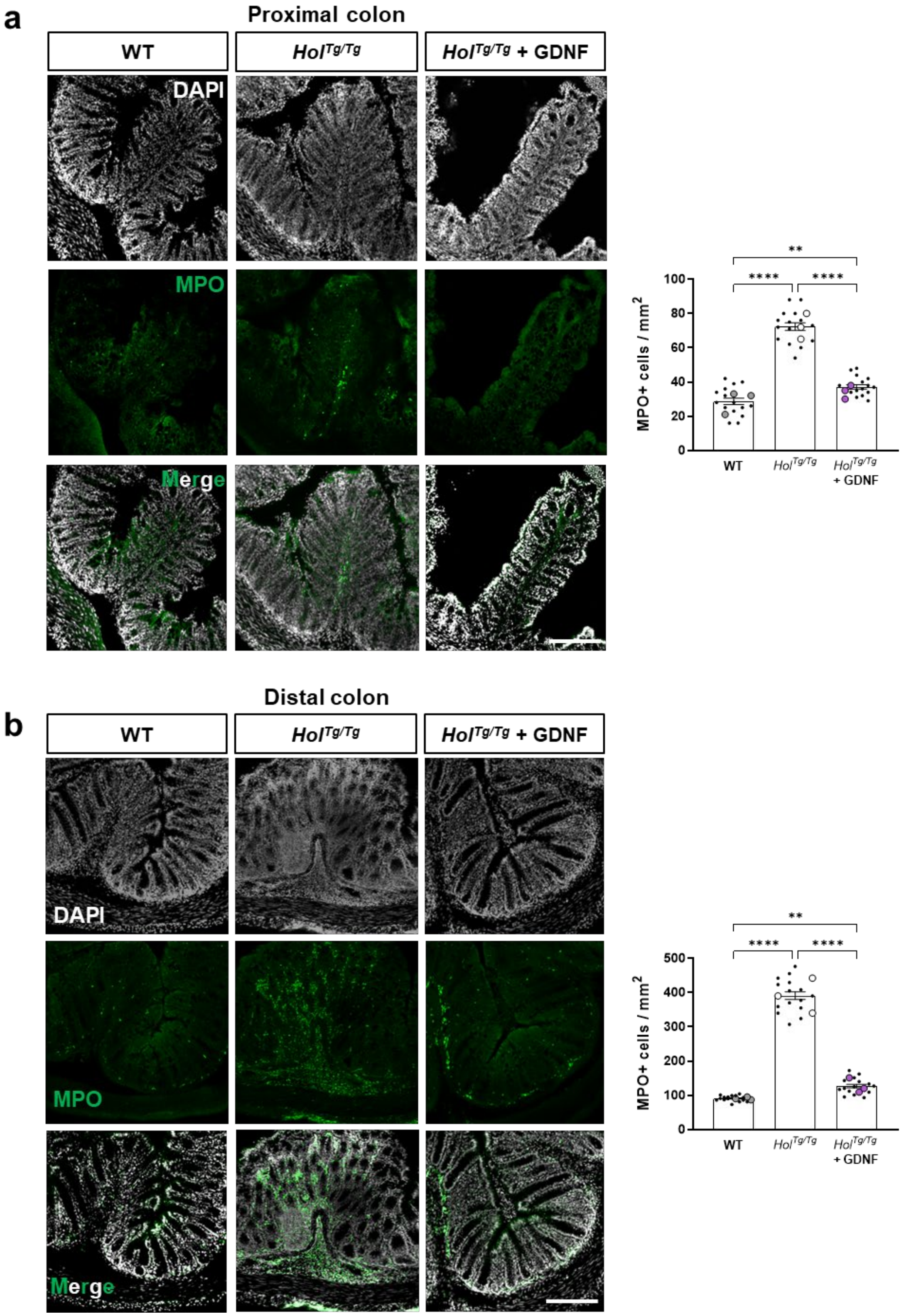


**Figure S3. GDNF treatment limits the infiltration of MPO$^+$ cells in both proximal and distal colon of P20 *Hol$^{Tg/Tg}$* mice.** (**a**,**b**) Representative images (left panels) and accompanying quantitative analysis (right panels) showing that GDNF treatment reduces MPO$^+$ cell density in both the proximal (a) and distal (b) colon of *Hol$^{Tg/Tg}$* mice. All images show a 15µm-thick z-stack projection (scale bar, 70 µm). Large dots in the quantitative analysis indicate the mean value for each mouse (n=3 mice per group), while small black dots indicate each microscopic field analyzed (5 fields of view per animal). **p < .01, ****p < .0001; one-way ANOVA with Tukey's post-hoc test.

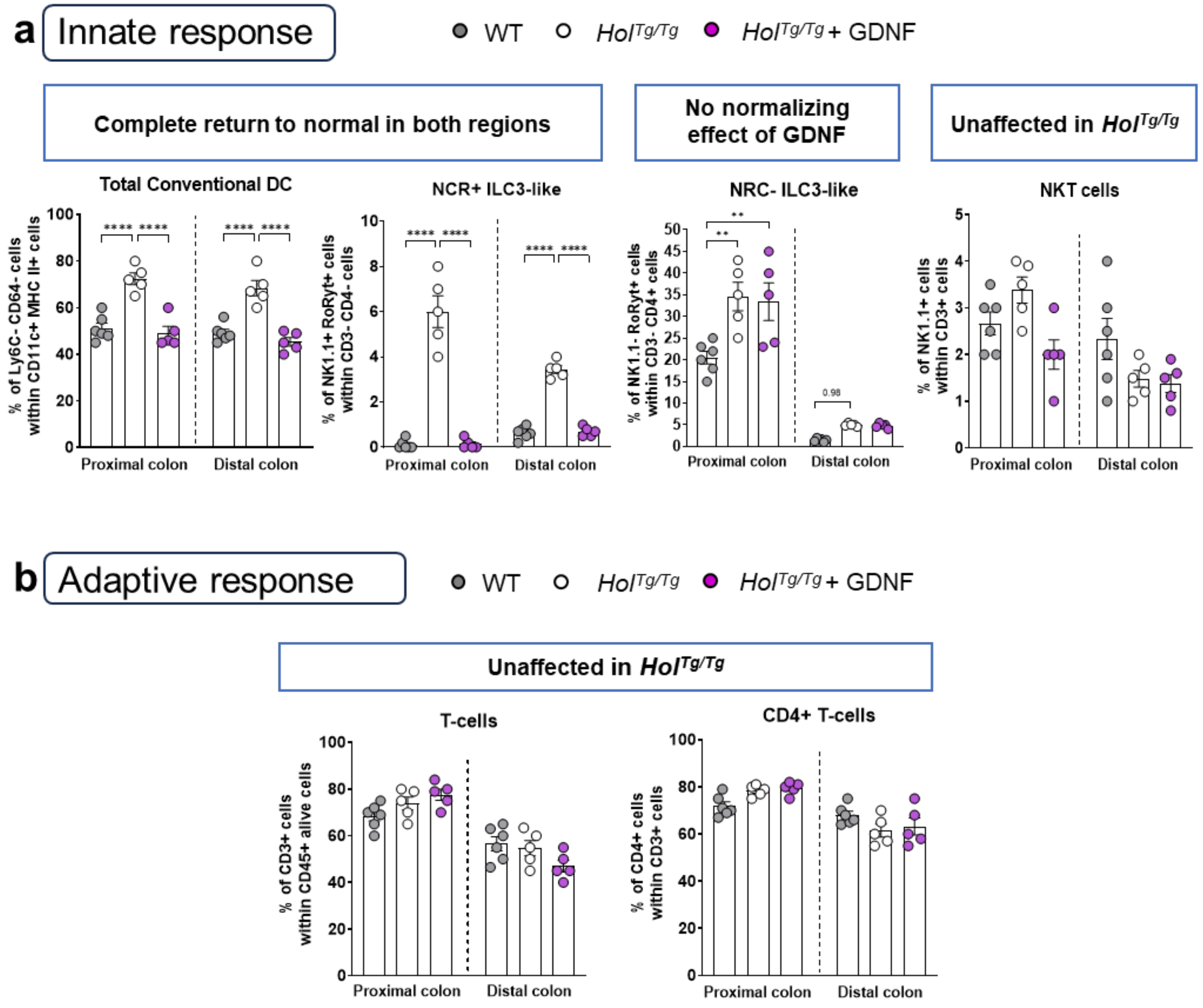


**Figure S4. Supplemental data showing that GDNF treatment influences both innate and adaptive immune responses in the colon of P20 *$Hol^{Tg/Tg}$* mice**. (**a**,**b**) Complementary graphs of the diverse innate (a) and adaptive (b) responses to GDNF treatment (n=5-6 mice per group). $^{**}p < .01$, $^{****}p < .0001$; two-way ANOVA with Sidak's post-hoc test. *In support of main Fig.4.*

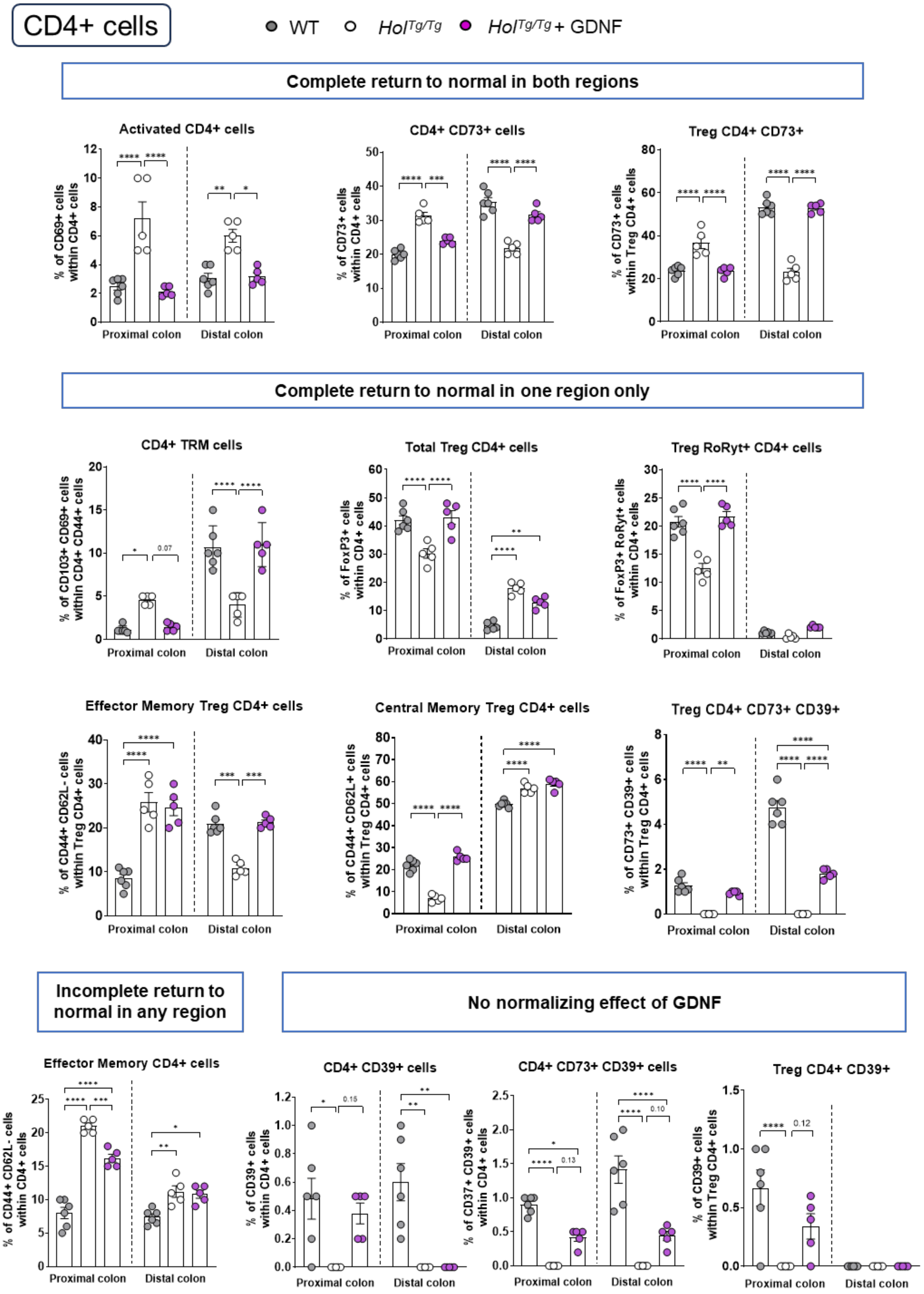


**Figure S5. Supplemental data showing that GDNF treatment impacts multiple CD4⁺ T-cell subsets in at least one colonic region of P20 *Hol*$^{Tg/Tg}$ mice**. Complementary graphs of the diverse responses of CD4⁺ T-cell subsets to GDNF treatment (n=5-6 mice per group). *p < .05, **p < .01, ***p < .001, ****p < .0001; two-way ANOVA with Sidak's post-hoc test. *In support of main Fig.5.*

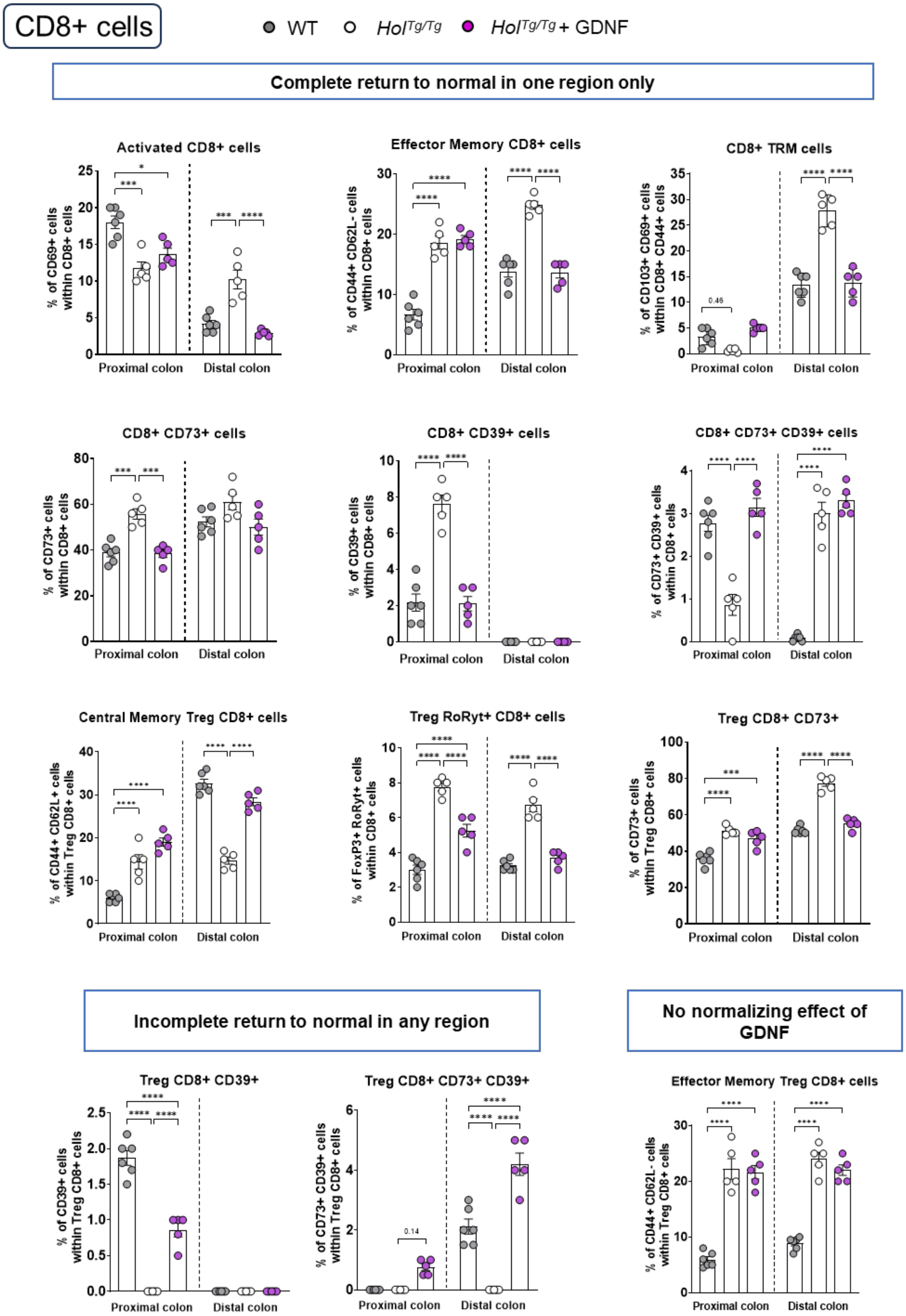


**Figure S6. Supplemental data showing that GDNF treatment impacts multiple CD8⁺ T-cell subsets in at least one colonic region of P20 *Hol*$^{Tg/Tg}$ mice**. Complementary graphs of the diverse responses of CD8⁺ T-cell subsets to GDNF treatment (n=5-6 mice per group). *p < .05, ***p < .001, ****p < .0001; two-way ANOVA with Sidak's post-hoc test. *In support of main Fig.6.*

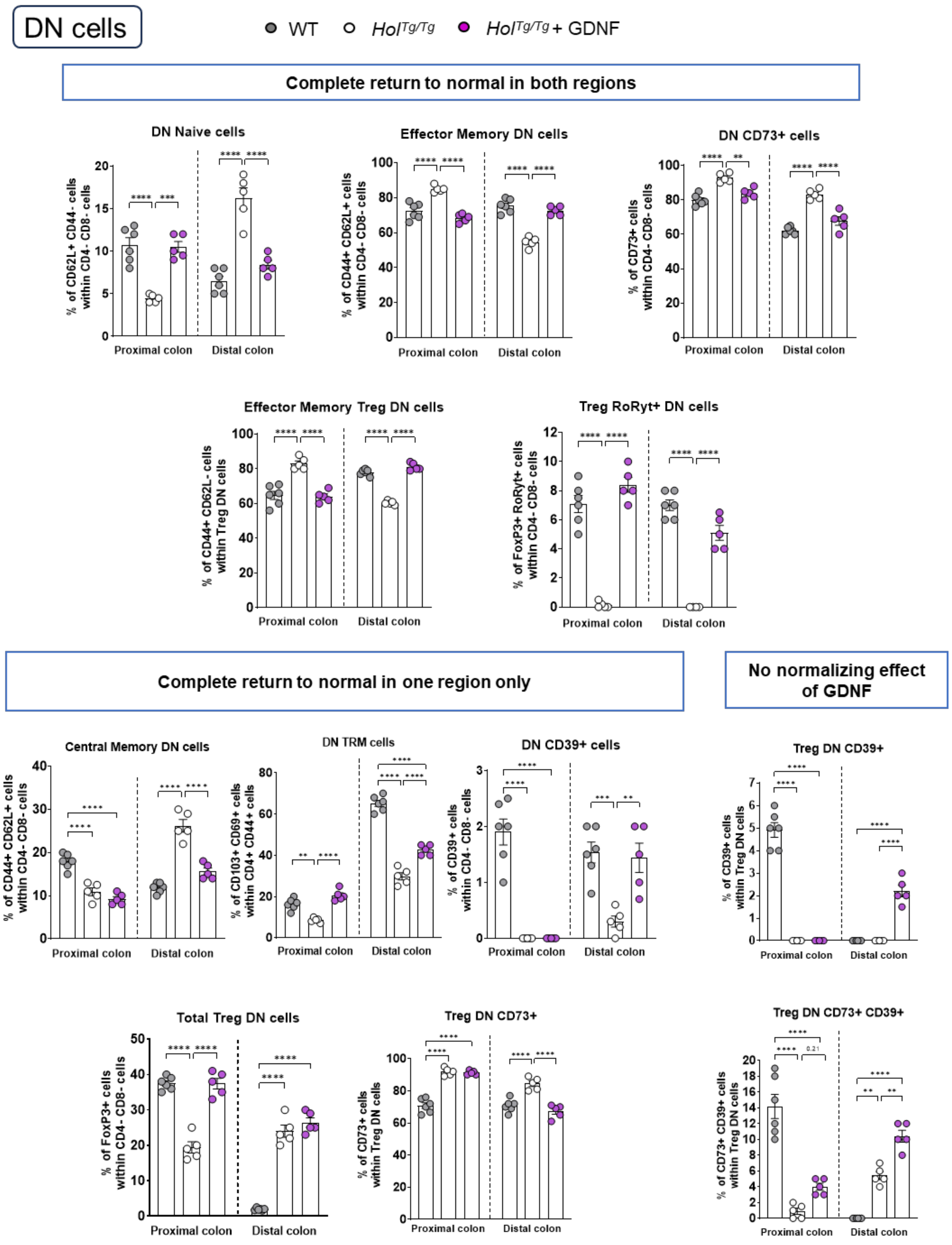


**Figure S7. Supplemental data showing that GDNF treatment impacts multiple CD4-CD8 double-negative T-cell subsets in at least one colonic region of P20 *Hol*$^{Tg/Tg}$ mice.** Complementary graphs of the diverse responses of CD4-CD8 double-negative T-cell subsets to GDNF treatment (n=5-6 mice per group). **p < .01, ***p < .001, ****p < .0001; two-way ANOVA with Sidak's post-hoc test. *In support of main Fig.7.*

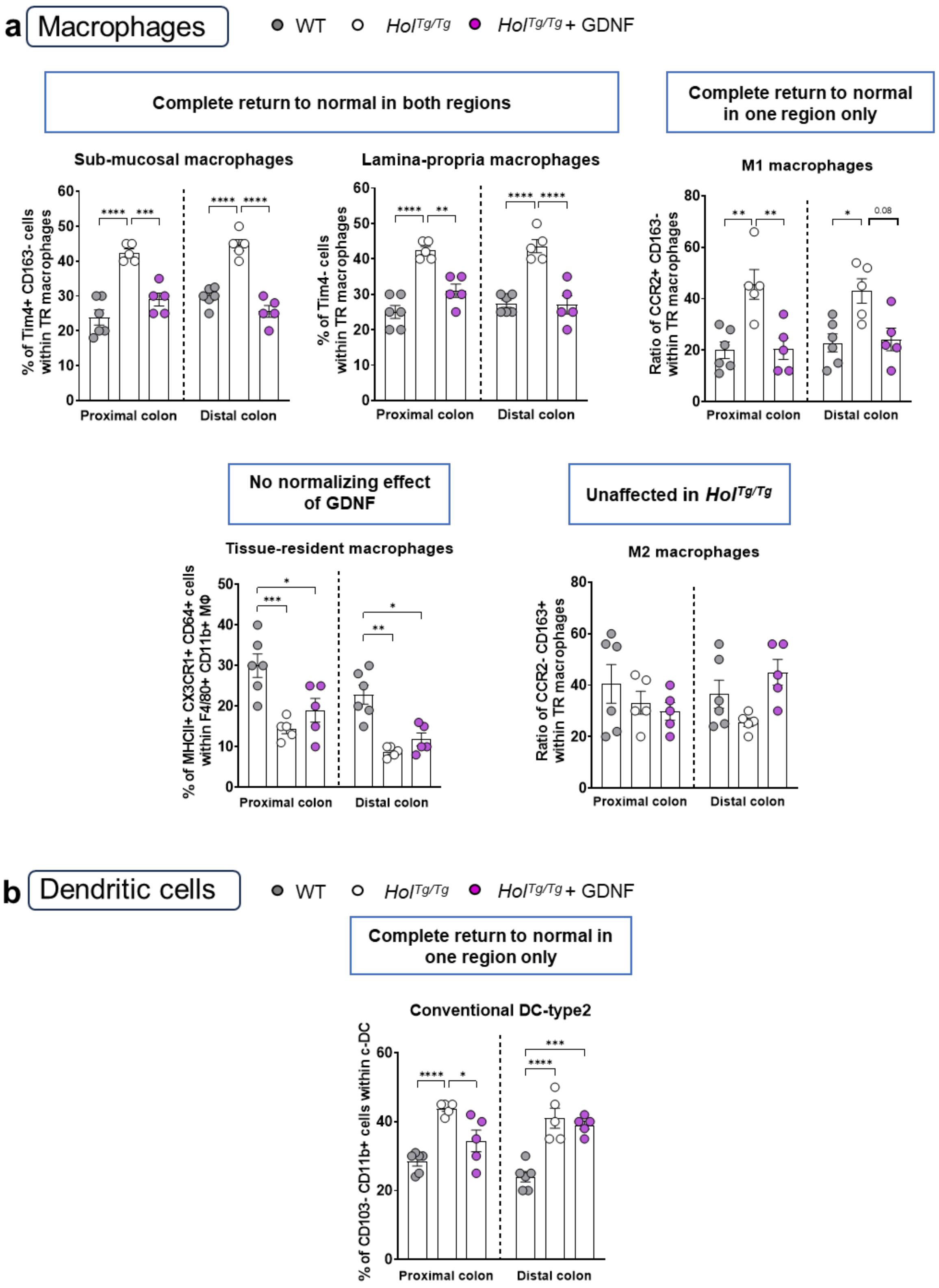


**Figure S8. Supplemental data showing that GDNF treatment impacts multiple macrophage and dendritic cell subsets in at least one colonic region of P20 *Hol*$^{Tg/Tg}$ mice**. Complementary graphs of the diverse responses of macrophage (a) and dendritic cell (b) subsets to GDNF treatment (n=5-6 mice per group). *p < .05, **p < .01, ***p < .001, ****p < .0001; two-way ANOVA with Sidak's post-hoc test. *In support of main Fig.8-9.*

**Table S1: List of antibodies and FISH probes utilized**

| Antibodies | | | | | | |
|---|---|---|---|---|---|---|
| | Target Protein | Purpose | Source | RRID | Host | Dilution |
| Primary Abs | ZO1 | Tight junctions | Invitrogen 33-9100 | AB_2533147 | Mouse | 1:500 |
| | CLDN3 | Tight junctions | Invitrogen 34-1700 | AB_2533158 | Rabbit | 1:200 |
| | DSG2 | Desmososme | Proteintech 21880-1-AP | AB_10836933 | Rabbit | 1:250 |
| | MPO | Neutrophils | Abcam Ab9535 | AB_307322 | Rabbit | 1:500 |
| | Target | Fluorophore | Source | RRID | Host | Dilution |
| Secondary Abs | Mouse | Alexa Fluor 647 | Jackson Immuno Research Labs 715-605-150 | AB_2340862 | Donkey | 1:500 |
| | Rabbit | Alexa Fluor 594 | Jackson Immuno Research Labs 711-585-152 | AB_2340621 | Donkey | 1:500 |
| | Mouse | Alexa Fluor 488 | Jackson Immuno Research Labs 715-545-150 | AB_2340846 | Donkey | 1:500 |

| FISH probes | | |
|---|---|---|
| **Probes** | Company | Sequences |
| **Eub338** | ThermoFisher (Custom oligonucleotides) | 5'-Cy3-GCTGCCTCCCGTAGGAGT-3' |
| **Non Eub (Negative control)** | | 5'-Cy3-CGACGGAGGGCATCCTCA-3' |
